# Whole genome duplication through mitotic slippage causes nuclear instability

**DOI:** 10.1101/2025.07.21.665898

**Authors:** S Gemble, M Budzyk, A Simon, R Lambuta, N Weiss, A Forest, YA Miroshnikova, F Scotto Di Carlo, V Marthiens, C Verdel, J Fang, C Desdouets, SA Wickström, G Ciriello, E Oricchio, G Almouzni, R Basto

## Abstract

Whole-genome duplication (WGD), leading to polyploidy can arise in physiological and pathological contexts^1–5^. WGD can occur via non-canonical cell cycles such as mitotic slippage, cytokinesis failure or endoreplication^1,3^. Whether the routes to WGD influence the behaviour of the resulting polyploid cells remains unclear. Here, we compared these routes under both physiological and non-physiological conditions. Remarkably, only mitotic slippage led to widespread nuclear abnormalities defined by highly variable nuclear deformations that we termed nuclear instability. Mechanistically, we found that these nuclei were softer - due to high levels of histone 3 phosphorylation in G1 altering chromatin compaction - and thus more vulnerable to microtubule-driven deformations. The resulting nuclear instability leads to local nuclear reorganisation and changes in 3D genome organisation impacting ultimately gene expression. Importantly, we observed similar nuclear instability in megakaryocytes, which are physiological polyploid cells that we show here to be generated by mitotic slippage, providing a molecular mechanism for their atypical nuclear architecture^6,7^. In striking contrast, nuclear shape was stable in different physiological polyploid cells generated by cytokinesis failure and endoreplication. Overall, our findings highlight that the route towards WGD matters and that mitotic slippage uniquely destabilizes nuclear architecture, with implications for both physiology and disease.

## Introduction

Most animal cells are diploid, carrying two copies of each chromosome (2N). Through a developmentally regulated and reproducible program, certain cells increase their chromosome number through whole genome duplication (WGD), accumulating multiple sets of each chromosome (> 2N) – hereafter called physiological polyploidy^8^. In most cases, the increase in DNA content is related with an increase in metabolic activity, barrier function or even regenerative capacity. For instance, in mammals, megakaryocytes become polyploid (modal ploidy 16N) to increase their metabolic capacity and produce a high number of platelets^7,9^ while salivary gland cells become polyploid (1024N) during *Drosophila* larval development to produce large quantities of glue, which is necessary for *Drosophila* pupation^8,10,11^. Mammalian hepatocytes also become polyploid (8N) during liver maturation. The role of polyploidy in this context is still debated, but previous studies suggested that polyploid hepatocytes may have higher metabolic and regenerative capacities^12,13^. In contrast, when WGD occurs in cells that should not become polyploid - hereafter called unscheduled polyploidy - it leads to genetic and chromosomal instability^2,14–20^. Furthermore, the resulting tetraploidy (4N) - that results from a single WGD - is a common feature of human tumours. Indeed, about 30% of tumours are tetraploid, reaching up to 70% in some specific cases like non-small cell lung carcinomas^4,5,18,21^.

The canonical cell cycle typical of somatic cells relies on coupling genome duplication with mitotic cell division to generate two diploid daughter cells^22^. Polyploid cells can be generated by alternative cell cycles - hereafter called non-canonical cell cycles. These cell cycles rely on incomplete or absent mitosis uncoupling the duplication of the genome from cell division leading thus to WGD^1^. Cells can exit mitosis before or after chromosome segregation through mitotic slippage (MS) or cytokinesis failure (CF), respectively (Fig. 1A). Otherwise, cells can alternate S-phase with G-phase using a cell cycle called endoreplication (EnR) (Fig. 1A). MS and EnR generate polyploid cells with a single nucleus, while CF generates binucleated polyploid cells^6,11,23^. However, the functional consequences of these non-canonical cell cycles remain poorly understood. Indeed, while MS, CF and EnR have been proposed to take place in physiological and pathological contexts^1,2,8,12,24^, it is still not clear whether different means to reach WGD lead to different outcomes.

**Figure 1:**
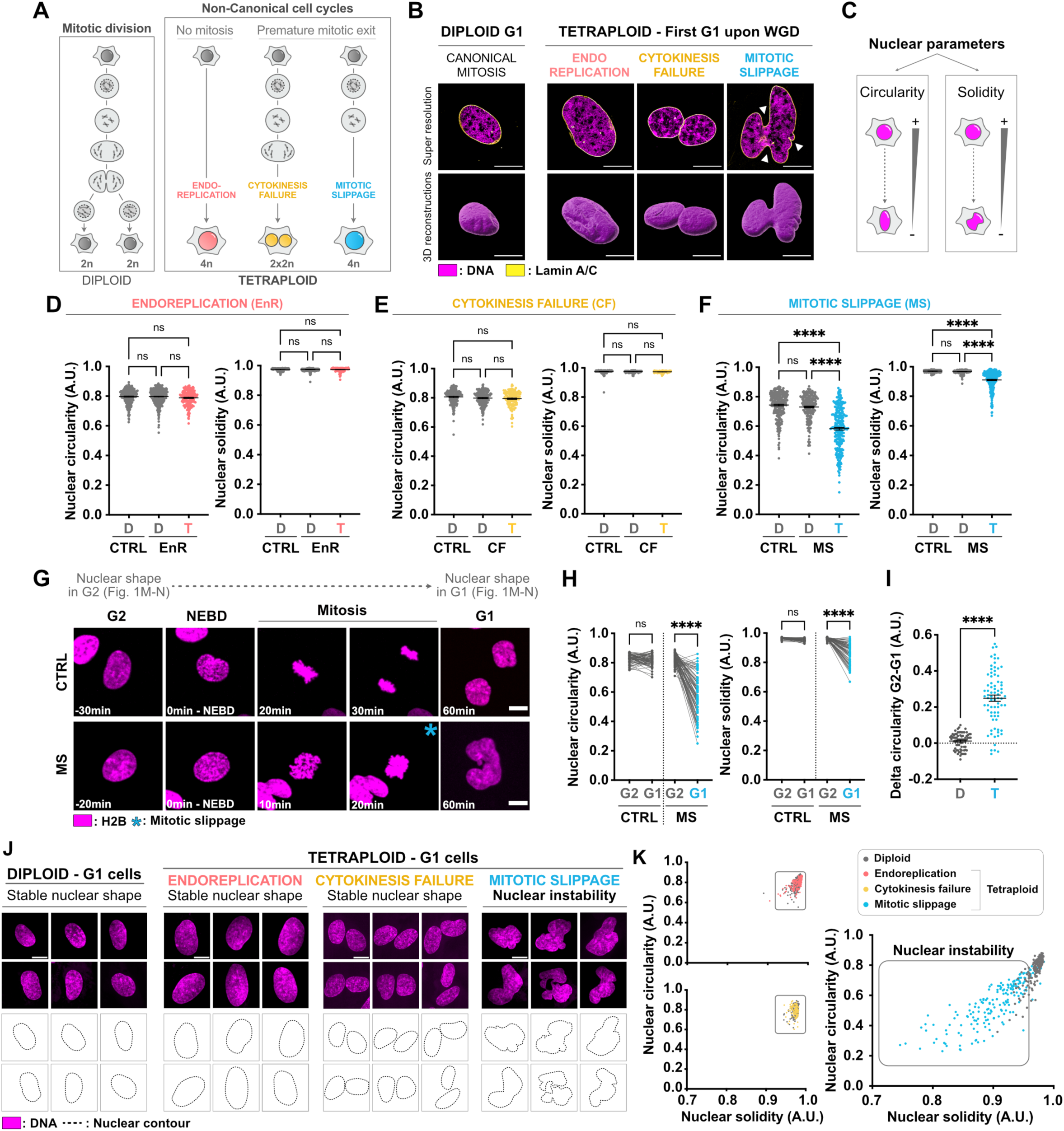
Unscheduled WGD induced through MS generates cells with nuclear instability. **(A)** Schematic representations of mitotic division (left panel) and of the different means employed to generate tetraploid cells (right panel). **(B)** Structural illumination microscopy (SIM) representative images (top) and their corresponding 3D reconstructions (bottom) showing nuclear shape in G1 RPE-1 cells upon canonical mitotic division, MS, CF or EnR. Lamin A/C in yellow, DNA in magenta. **(C)** Schematic illustration of the circularity and solidity indices. **(D-F)** Graphs showing the nuclear circularity and solidity indices in RPE-1 G1 cells upon **(D)** EnR (in red), **(E)** CF (in yellow) and **(F)** MS (in blue). Mean ± SEM, >150 G1 cells analysed per condition, three independent experiments. **(G)** Stills from time-lapse imaging of RPE-1 cells stably expressing mCherry-H2B (in magenta) performing mitotic division (upper panel) or MS (lower panel). **(H)** Graph presenting the nuclear circularity and solidity measured at the single cell level in the same cells in G2 and in G1 upon mitotic division (in grey) or MS (in blue). >80 cells analysed per condition, three independent experiments. **(I)** Graph showing the delta circularity (defined as the difference between nuclear circularity in G2 vs in G1) upon mitotic division (in grey) or MS (in blue) in RPE-1 cells. Mean ± SEM, >75 cells analysed per condition, three independent experiments. **(J)** Representative images showing G1 RPE-1 nuclei upon mitotic division, MS, CF and EnR (upper panel). DNA in magenta. The nuclear contour is indicated (black dotted line, lower panel). **(K)** Graph showing the nuclear circularity and solidity index from **(D-F)** in diploid (in grey) and in tetraploid G1 RPE-1 cells generated by EnR (in red), CF (in yellow) and MS (in blue). Scale bars: 10µm. WGD=whole genome duplication. D=diploid; T=tetraploid; MS=mitotic slippage; CF=cytokinesis failure; EnR= endoreplication. NEBD=nuclear envelope breakdown. A.U.=arbitrary unit. **(D,E,F)** ANOVA test (one-sided). **(H,I)** t-test (two-sided). ns=not significant. *=P ≤ 0.05. **= P ≤ 0.01. ***= P ≤ 0.001. ****= P ≤ 0.0001.

Here, by combining comparative analyses integrating physiological and unscheduled models of WGD, we found that MS, unlike CF and EnR, generates polyploid cells with irregular nuclear architecture. Mechanistically, we identified that MS leads to mitotic exit with high levels of histone 3 phosphorylation on serine 10 (H3S10). This impairs chromatin compaction compromising nuclear stiffness and allowing microtubules to promote nuclear deformability. This induces local nuclear reorganisation and changes in 3D genome organization and in gene expression. Importantly, we observed that a similar mechanism contributes to the irregular nuclear shape of megakaryocytes, physiological polyploid cells generated by MS. Overall, these data highlight the fact that different means to generate WGD have different outcomes, which may influence downstream cell functions.

## Results

### Mitotic slippage, unlike cytokinesis failure and endoreplication, generates tetraploid cells with nuclear deformations

To define whether different routes of WGD have different outcomes, we induced unscheduled WGD through mitotic slippage (MS), cytokinesis failure (CF) or endoreplication (EnR) in RPE-1 cells (see methods, Fig. 1A and Supp. Fig. 1A)^15,25^, which are p53-proficient diploid human cell line. As previously described and due to the methods employed to induce WGD^15^, this strategy generates a mixed population of diploid (2N) and tetraploid (4N) cells that can be distinguished by nuclear and cell size^15^. Importantly, to avoid any effect due to cell cycle progression, cells were synchronised in the first G1 upon WGD using the CDK4/6 inhibitor Palbociclib^26^ at 1µM.

Using these highly comparable systems, we observed that MS, but not CF and EnR, generated tetraploid cells with abnormal nuclear shape (Fig. 1B). To quantitatively analyse the nuclear architecture in these cells, we measured nuclear circularity and solidity, two indices, informing on nuclear elongation and on the presence of nuclear invaginations, respectively (Fig. 1C). We observed that tetraploid cells generated through MS showed decreased nuclear circularity and solidity, while CF and EnR showed values similar to diploid cells (Fig. 1D-F). Similar results were obtained using alternative methods and other diploid cell lines - BJ and HCT116 (Supp. Fig. 1A-H). Additionally, when cells were cultured in 3D, nuclear deformations upon MS were still detected (Supp. Fig. 1I-L). Inducing MS in asynchronous cells, also showed similar results (Supp. Fig. 1M). Interestingly, we found that the nuclear area was similar in MS- and EnR-generated newly born tetraploid cells (Supp. Fig. 1N-O), suggesting that nuclear defects upon MS does not affect the overall nuclear size but only nuclear architecture.

We next performed live imaging approaches using cells stably expressing mCherry-H2B, to follow the transition from G2 to the subsequent G1 at the single cell level. In control diploid cells that undergo canonical mitotic divisions, nuclear circularity and solidity from G2 to G1 remained stable (Fig. 1G-H and Movie 1). In contrast, in tetraploid cells generated by MS an obvious decrease could be noticed (Fig. 1G-H and Movie 2). Plotting the delta circularity (defined as the difference between nuclear circularity in G2 vs in G1) showed values close to 0 in diploid cells (Fig. 1I). This indicates that nuclear architecture was similar before and after mitosis in diploid cells - i.e. nuclear architecture was stable over subsequent cell cycles, as expected. Delta circularity increase in MS-induced tetraploid cells indicates that MS impairs the re-establishment of a typical nuclear architecture in newly born tetraploid cells. Indeed, we observed that MS-induced tetraploid G1 cells exhibited heterogenous nuclear shapes (Fig. 1J), that were still present in the following G2 (Supp Fig. 1P). To understand whether these abnormal shapes were maintained over multiple cell cycles, we induced MS in a p53-depleted RPE-1 cell line, which is permissive to tetraploid cell proliferation^15,27–29^. Nuclear circularity and solidity remained low over multiple cell cycles in MS-generated tetraploid cells (Supp Fig. 1Q-S).

At the population level, we were struck by the level of heterogenous nuclear shapes upon MS. Indeed, we found that MS generated a population of tetraploid cells displaying a wide-range of nuclear circularity and solidity indexes (Fig. 1J-K). This reflects that WGD when induced through MS generates a high degree of nuclear shape diversity that seems stochastic - a phenomenon that we called here nuclear instability. In contrast, diploid, CF- and EnR-generated tetraploid cells are clustered in a homogenous cell population with nuclear circularity and solidity indexes close to 1, indicating that these cells have a uniform nuclear architecture (Fig. 1J-K).

Together, these findings show that MS generates nuclear instability - i.e. a high degree of nuclear shape diversity, while CF and EnR induction do not impair nuclear morphology.

### Upon mitotic slippage, cells enter G1 with high H3S10 phosphorylation levels

We then investigated why only MS generated nuclear instability. During mitotic entry, phosphorylation of Ser10 of Histone 3 (H3S10) promotes nuclear envelope breakdown^30–32^ (Supp. Fig. 2A-B). As cells exited mitosis, the phosphorylation levels gradually decreased allowing nuclear envelope reformation^30,31^ (Supp. Fig. 2A-B). Since EnR cells do not enter mitosis, H3S10 phosphorylation levels should remain low throughout the cell cycle, while CF cells exit mitosis in a state in which H3S10 phosphorylation levels should also be low^30,31^. In contrast, MS cells, that exit mitosis earlier, should display high H3S10 phosphorylation levels when exiting mitosis^30,31^. In line with this model, we found that MS-induced tetraploid cells still have H3S10 phosphorylation in the first G1 upon WGD - forming patches close to the nuclear envelope (Fig. 2A-B). In contrast, diploid or CF-induced tetraploid cells do not exhibit H3S10 phosphorylation in G1 (Fig. 2A-B), as expected^30,31^.

**Figure 2:**
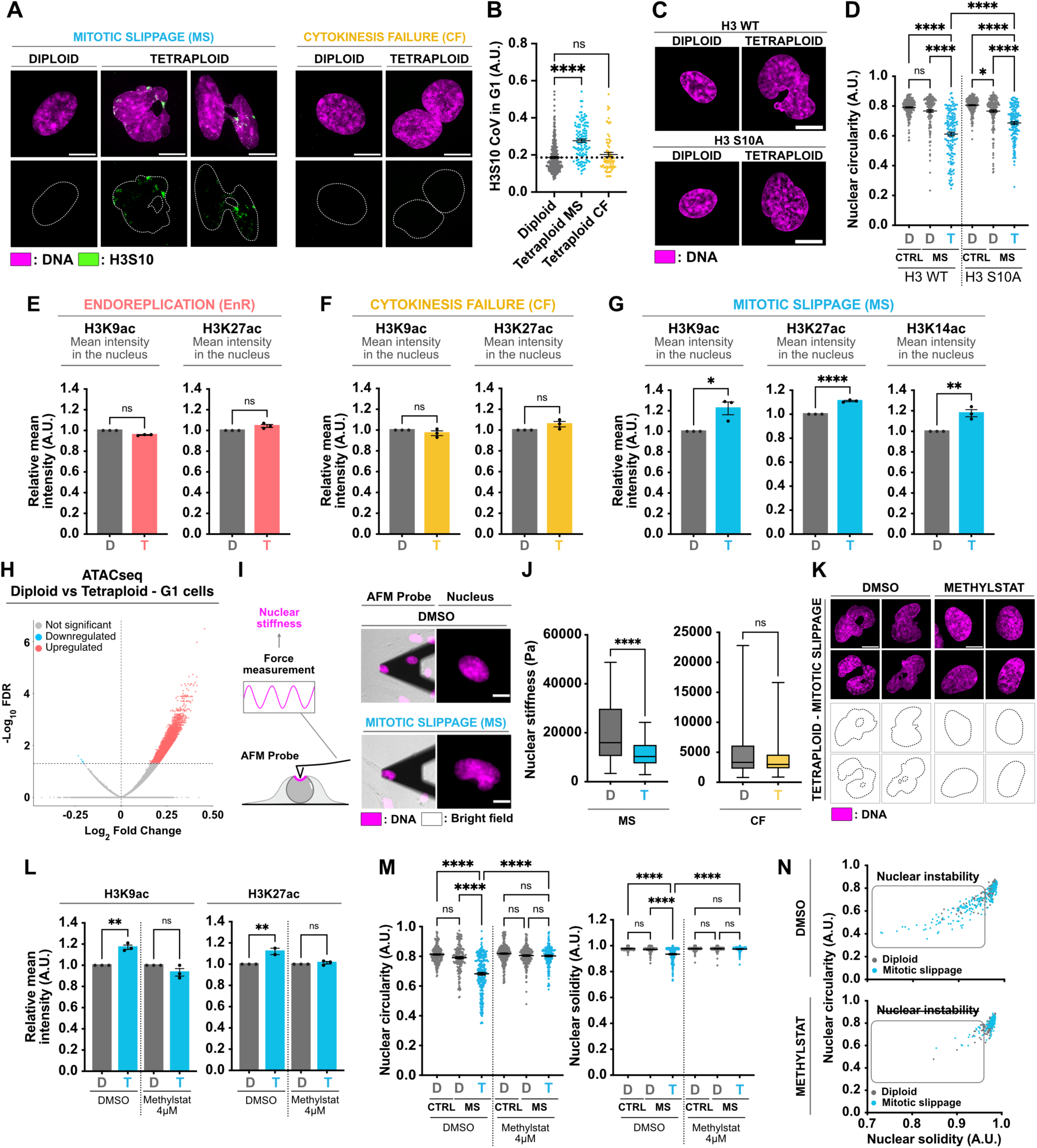
Mitotic slippage generates tetraploid cells with high histone acetylation levels and softer nuclei promoting nuclear deformations **(A)** SIM representative images showing diploid and tetraploid G1 RPE-1 cells generated by MS or EnR. H3S10 in green, DNA in magenta. **(B)** Graph showing the H3S10 coefficient of variation (defined as the ratio of the standard deviation to the mean) in G1 in diploid (in grey) or tetraploid RPE-1 cells generated by MS (in blue) or by CF (in yellow). Mean ± SEM, >75 cells analysed per condition, three independent experiments. **(C)** Representative images showing diploid and tetraploid G1 RPE-1 cells generated by MS and stably expressing H3 WT or H3 S10A. DNA in magenta. **(D)** Graph showing the nuclear circularity index in diploid (in grey) and in MS-generated tetraploid (in blue) G1 RPE-1 cells stably expressing H3WT or H3S10A. Mean ± SEM, >150 cells analysed per condition, three independent experiments. **(E-G)** Graphs showing H3K9ac, H3K27ac and H3K14ac levels in diploid (in grey) and in tetraploid G1 RPE-1 cells generated by EnR (**E**, in red), by CF (**F**, in yellow) or by MS (**G**, in blue). Mean ± SEM, >100 cells analysed per condition, three independent experiments. **(H)** Volcano plot showing ATACseq data comparing diploid and tetraploid G1 RPE-1 cells generated by MS. **(I)** Left panel - Schematic representation showing the principle of atomic force microscopy (AFM). Right panel - Representative images showing diploid and MS-generated tetraploid G1 RPE-1 cells. DNA in magenta. Brightfield in grey. **(J)** Graph showing nuclear stiffness in diploid (in grey) and in tetraploid G1 RPE-1 cells generated by MS (in blue) or by CF (in yellow). Box and whiskers ± Min to Max, >90 cells (left panel) or >65 cells (right panel) analysed per condition, three independent experiments. **(K)** Representative images showing diploid and tetraploid G1 RPE-1 cells generated by MS treated with DMSO or Methylstat. DNA in magenta. **(L)** Graph showing H3K9ac and H3K27ac levels in diploid (in grey) and in MS-generated tetraploid (in blue) G1 RPE-1 cells treated with DMSO or Methylstat. Mean ± SEM, >100 cells analysed per condition, three independent experiments. **(M)** Graph showing the nuclear circularity and solidity indices in diploid (in grey) and in MS-generated tetraploid (in blue) G1 RPE-1 cells treated with DMSO or Methylstat. Mean ± SEM, >125 cells analysed per condition, three independent experiments. **(N)** Graph showing the nuclear circularity and solidity indices from **(M)** in diploid (in grey) and tetraploid G1 RPE-1 cells generated by MS (in blue) treated with DMSO or Methylstat. Scale bars: 10µm. D=diploid; T=tetraploid; MS=mitotic slippage. EnR=endoreplication. CF=cytokinesis failure. A.U.=arbitrary unit. **(J)** Kolmogorov-Smirnov test. **(E,F,G,L)** t-test (two-sided). **(B,D,M)** ANOVA test (one-sided). ns=not significant. *=P ≤ 0.05. **= P ≤ 0.01. ***= P ≤ 0.001. ****= P ≤ 0.0001.

We next wanted to directly test the contribution of high H3S10 phosphorylation levels to nuclear architecture. We generated a cell line, expressing an H3 mutant version, where Serine 10 was replaced by an Alanine residue - H3S10A^30^, in the background of wild type (WT) H3. A partial decrease in H3S10 phosphorylation during mitosis was noticed as expected (Supp. Fig. 2C). Importantly, this was sufficient to increase both nuclear circularity and solidity in newly born tetraploid MS-induced cells, improving their overall nuclear shape and thus decreasing nuclear instability (Fig. 2C-D and Supp. Fig 2D-E). To confirm that maintaining H3S10 phosphorylation in G1 was sufficient to generate nuclear deformations, we inhibited the phosphatase responsible for H3S10 dephosphorylation at mitotic exit using Calyculin A^33^ in diploid cells. This treatment resulted in increased H3S10 phosphorylation levels in diploid G1 cells and remarkably, nuclear deformations characterized by invaginations similar to those found in MS-generated tetraploid cells were noticed (Supp. Fig. 2F-G).

Together, our results show that low H3S10 phosphorylation at mitotic exit is a pre-requisite to re-establish proper nuclear morphology during nuclear re-assembly.

### Mitotic slippage generates tetraploid cells with high histone acetylation levels and low nuclear stiffness

We then investigated the mechanism underlying nuclear deformations upon MS. Previous work identified that H3S10 phosphorylation impacts other histone modifications^34–37^. We thus quantified histone acetylation and methylation levels using structural illumination microscopy. The fluorescence intensity levels were measured in diploid and tetraploid G1 cells at the level of the whole nucleus (Supp. Fig. 3A).

We first noticed a global increase in H3K9ac, H3K27ac and H3K14ac levels in tetraploid G1 cells generated by MS (Fig. 2E-G and Supp. Fig. 3B). In contrast, in CF-or EnR-induced tetraploid G1 cells, histone acetylation remained similar to diploid cells (Fig. 2F-G and Supp. Fig. 3B). Importantly, reducing H3S10 phosphorylation by expressing H3S10A mutant was sufficient to prevent the increase in acetylation levels observed upon MS (Supp. Fig. 3C). This suggests that high H3S10 phosphorylation levels upon MS induced a global increase in histone acetylation in tetraploid G1 cells.

Since histone acetylation is commonly associated with open chromatin/euchromatin^38^, we performed ATACseq and found that chromatin accessibility was higher in MS-induced tetraploid G1 cells compared to diploid cells (Fig. 2H).

When measuring histone methylation, usually associated with closed chromatin/heterochromatin^39,40^, we found that only H3K9me2 levels were increased upon MS, while H3K9me3 and H3K27me3 levels remained similar to diploid cells (Supp. Fig 3D-F). In CF- or EnR-generated cells, no changes in histone methylation levels were found (Supp. Fig. 3G-H). Interestingly, we also observed that MS led to a decrease in the number of Heterochromatin Protein 1⍺ (HP1⍺) foci (Supp. Fig. 3I-J), a key component known to promote nuclear compaction^41^. Together, these findings demonstrate that MS impairs histone compaction in newly born tetraploid cells favouring an overall open chromatin.

It is well established that several factors, including chromatin states, contribute to nuclear envelope deformability counteracting cytoplasmic forces that act on the nucleus^42,43^. It is therefore possible that the nuclear deformations of tetraploid cells induced through MS reflected a decrease in nuclear mechanical properties due to changes in chromatin compaction as described above. To test this possibility, we first tested whether MS alters nuclear stiffness using atomic force microscopy (AFM) (Fig 2I). Using two different strategies to generate MS, we found that G1 tetraploid nuclei were significantly softer than their diploid counterparts (Fig. 2J and Supp. Fig. 3K). In contrast, CF-induced tetraploid G1 cells exhibit nuclear stiffness similar to diploid cells (Fig. 2J).

We next investigated the functional relationship between higher acetylation levels, decreased nuclear stiffness and nuclear deformations upon MS. We used two complementary strategies to decrease acetylation levels in MS-induced tetraploid cells. We treated cells with (1) C646, a histone acetylase inhibitor^44^, which decreases histone acetylation levels (Supp. Fig. 4A) or with (2) Methylstat, a histone demethylase inhibitor^45^, which promotes histone methylation preventing histone acetylation on the same residues (Supp. Fig. 4A-B). These treatments were sufficient to decrease histone acetylation and to promote nuclear compaction and thus nuclear stiffness preventing nuclear deformations and nuclear instability in MS-induced tetraploid G1 cells (Fig. 2K-N and Supp. Fig. 4A-F). These results were confirmed using JIB, another histone demethylase inhibitor^46^, by live imaging, using another method to induce MS and in BJ and HCT116 cells (Supp. Fig. 4G-O).

These data show that MS generates tetraploid cells with high levels of chromatin acetylation that contribute to decrease their nuclear stiffness.

### Microtubules generate nuclear invaginations in MS-induced tetraploid G1 cells

Detailed analysis of mitotic exit in diploid and MS-induced tetraploid cells revealed an increase in nuclear area, most likely due to chromosome decondensation^47^. However, in tetraploid cells, this was accompanied by the appearance of nuclear invaginations (Supp. Fig 5A, pink arrow). Indeed, while quantifying nuclear circularity and solidity, we found that these indices were quite stable in diploid cells during early G1 (Supp. Fig 5B, upper panel), whereas both parameters decreased in newly born tetraploid cells during mitotic exit (Supp. Fig 5B, lower panel).

Cytoskeleton components interact with the nuclear envelope, a process that contributes to the establishment of nuclear architecture^48^. Using time-lapse microscopy in cells incubated with SPY-Tubulin and SPY-Actin, we followed microtubules and actin, respectively. Characterization of cytoskeleton proteins in G1 cells showed that microtubules (but not actin) accumulated at nuclear invaginations in MS-induced tetraploid cells (Supp. Fig 5C-E), which was not seen in EnR-generated tetraploid cells (Supp. Fig. 5F). We found that just after nuclear envelope reformation upon MS, microtubules accumulated at invagination sites assembling in thick bundles whereas actin did not (Supp. Fig. 5G-H and Movies 3-4). These data suggest that microtubules unlike actin, contribute to nuclear instability in newly born tetraploid cells.

To functionally test the contribution of microtubules in generating nuclear invaginations, we removed microtubules using nocodazole, a microtubule-depolymerizing agent^49^, just before MS (Supp. Fig. 6A-B and Movie 5-6). Strikingly, in the absence of microtubules, the nuclear shape in newly born tetraploid cells became more regular, presenting fewer deformations in G1, as also revealed by a decrease in the delta circularity (Supp. Fig. 6B). Moreover, treating tetraploid cells with nocodazole was sufficient to enhance both nuclear circularity and solidity reducing thus nuclear instability (Supp. Fig. 6C-D). These results were confirmed using another mean to induce MS (Supp. Fig. 6E-G and Movie 7-8). We then tested whether stabilizing microtubules could exacerbate nuclear deformations. To do so, we treated tetraploid cells with low-doses of Taxol, a microtubule-stabilizing agent^50^, before MS. In these conditions, nuclear deformations were even more impressive and a decrease in both nuclear circularity and solidity was noticed associated with a higher nuclear instability (Supp. Fig. 6H-I).

We conclude that the microtubule cytoskeleton is the main contributor to nuclear deformations in newly born tetraploid cells induced by unscheduled MS.

### Nuclear deformations upon mitotic slippage leads to local nuclear reorganisation

We then investigated whether the nuclear deformations upon MS have a local impact on nuclear architecture. To do so, we conducted a mini screen in which 8 chromatin-related and 6 nuclear envelope-related candidates were visually examined using structural illumination microscopy. The candidate list was compiled based on their well-established roles in nuclear morphology^30,42,43,51–54^ and the fluorescence intensity levels were measured in diploid and tetraploid G1 cells at intact and deformed nuclear envelope regions.

This mini screen identified that H3K9me2 levels were decreased in specific regions that coincide with nuclear deformations in MS-induced tetraploid cells (Fig. 3A-D). Because the tools used to characterize H3K9me2 are the subject of controversial studies^55^, we also used another H3K9me2 antibody and two different protocols and observed the same results (Supp. Fig. 7A-C). H3K9me2 is normally enriched at the nuclear periphery and associated with chromatin regions that are less dynamic^30,54^. H3K9me2 coefficient of variation (CoV) at the nuclear envelope (defined as the ratio of the standard deviation to the mean) was increased in tetraploid cells generated by MS (Supp. Fig. 7D), indicating uneven distribution of this modification, in sharp contrast with the uniform distribution of H3K9me2 at the nuclear envelope of diploid nuclei (Fig. 3D and Supp. Fig. 7D). Similar defects in H3K9me2 localization at invaginations were found in tetraploid cells where MS was induced by a different mean, and in BJ and HCT116 cell lines (Supp. Fig. 7E-G). In contrast, we observed that both CF- and EnR-induced tetraploid cells displayed homogeneous distributions of H3K9me2 (Fig. 3A-C and Supp. Fig. 7H-I). Moreover, no local changes in H3K9me3, H3K27me3, H3K9ac and H3K27ac levels, were noticed upon MS (Supp. Fig. 7J-K).

**Figure 3:**
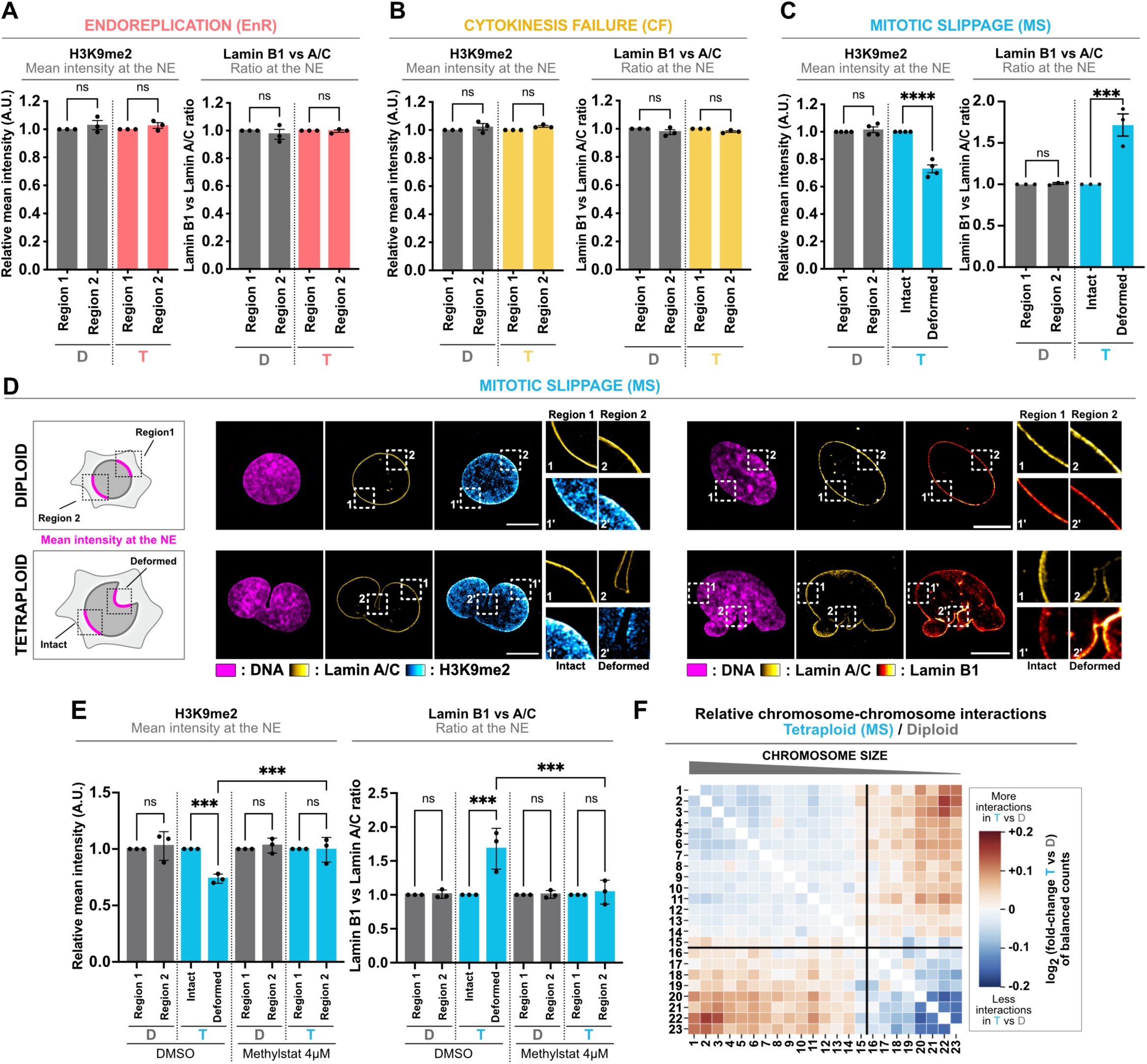
Mitotic slippage promotes local nuclear reorganisation and impairs 3D genome organisation **(A-C)** Graphs showing the relative H3K9me2 (left panel) or Lamin B1 vs A/C ratio (right panel) mean intensity at the nuclear envelope in diploid (in grey) and tetraploid G1 RPE-1 cells generated by EnR (**A**, in red), by CF (**B**, in yellow) or by MS (**C**, in blue). Mean ± SEM, >100 cells analysed per condition, three independent experiments. **(D)** Left, schematic representation fluorescence intensity measured at the nuclear envelope or locally at two random regions (in diploid cells) and at intact and deformed regions (in tetraploid cells). Middle and right panels, SIM representative images showing diploid (top) and tetraploid (bottom) G1 RPE-1 cells generated by MS. Lamin A/C in yellow hot, H3K9me2 in cyan hot, Lamin B1 in red hot. DNA in magenta. **(E)** Graphs showing the relative H3K9me2 or Lamin B1 vs A/C ratio mean intensity at the nuclear envelope in diploid (in grey) and in tetraploid G1 RPE-1 cells generated by MS (in blue) treated with DMSO or Methylstat. Mean ± SEM, >100 cells analysed per condition, three independent experiments. **(F)** Heatmap of the ratios of interchromosomal contact enrichments (observed versus expected) between tetraploid (MS) and diploid G1 RPE-1 cells. Chromosomes were sorted by length. Scale bar: 10µm. D=diploid. T=tetraploid. MS=mitotic slippage. EnR=endoreplication. CF=cytokinesis failure. NE= nuclear envelope. A.U.=arbitrary unit. (**A,B,C,E**) t-test (two-sided). ns=not significant. *=P ≤ 0.05. **= P ≤ 0.01. ***= P ≤ 0.001. ****= P ≤ 0.0001.

The Lamina, composed of a specific ratio of Lamin A/C and Lamin B proteins, plays a crucial role in maintaining nuclear architecture^43,51,52,56,57^. The ratio between Lamin A/C and B is crucial in establishing nuclear morphology^51,52,56,57^. Indeed, nuclear resistance is strongly impacted by Lamin A/C and a ratio dominated by Lamin B has thus been associated with soft tissues^56,57^. By measuring this ratio, we found differences in tetraploid cells at nuclear deformations regions. Frequently, Lamin B1 dominated the lamina network in invaginated domains, while the levels of Lamin A/C were lower than the ones found in intact regions (Fig. 3A-D). In line with this, we found that Lamin B Receptor (LBR) levels - a protein associated with soft tissues^58^ - were also increased locally at the nuclear deformations upon MS, while we found no changes in Lap2**β**, Emerin and Nuclear pore complex levels (Supp Fig. 7L). Similar defects were observed in MS-generated tetraploid cells using an alternative method and in BJ and in HCT116 cell lines (Supp. Fig. 7E-G). In contrast, we observed that both EnR- and CF-induced tetraploid cells displayed homogeneous Lamin ratios (Fig. 3A-C and Supp. Fig. 7H). Importantly, preventing nuclear deformations upon MS by reducing histone acetylation or by depolymerizing microtubules was sufficient to restore homogeneous distributions of H3K9me2 and Lamin ratios in tetraploid cells (Fig. 3E and Supp. Fig. 7M).

Together, our findings show that nuclear deformations upon MS promote local changes in nuclear organisation.

### Mitotic slippage impairs 3D genome organisation and gene expression

We next investigated the functional consequences of nuclear deformations observed upon MS. Recently it has been shown that WGD in p53-deficient cells leads to loss of chromatin segregation explained by an increased proportion of contacts between chromosomes, compartments, and chromatin domains^59^. Insulation of topology associated domains (TADs) requires a certain number of proteins that do not seem to scale with the tetraploid genome^59^. We wanted to investigate whether such defects were already detectable in the first G1 after MS, in the presence of p53. To do so, we isolated diploid and tetraploid cells using flow cytometry, as previously described^15^ (Supp. Fig. 8A-B). Importantly, cell sorting did not affect nuclear deformations in MS-generated tetraploid cells (Supp. Fig. 8C). Using high throughput chromatin capture conformation (HiC), we compared interactions at the chromosomal level and between chromatin domains in diploid and newly born MS-generated tetraploid G1 cells. Similarly to what has been previously observed, we found an increased number of contacts between short and long chromosomes (Fig. 3H) and between A and B compartments (Supp. Fig. 8D-G) in tetraploid cells compared to diploid cells.

Our findings showing changes in genome organisation and chromatin state / accessibility upon MS led us to test whether MS alters gene expression. We thus analysed newly born tetraploid cells in the first G1 upon MS by RNAseq. We found that 150 genes were deregulated in tetraploid G1 cells compare to diploid G1 cells (Supp. Fig. 8H).

Overall, our results identified that MS perturbs 3D genome organisation in newly born tetraploid cells culminating with changes in gene expression.

### Megakaryocytes undergo mitotic slippage and display nuclear instability

We decided to determine whether physiological polyploid cells generated by MS also exhibit nuclear deformations. To do so, we analysed megakaryocytes, differentiated polyploid cells, derived from hematopoietic progenitors^7^, and responsible for production of RNAs and proteins that will be transported to the cell tips to generate anucleate platelets^9^. To obtain megakaryocytes, we isolated mouse diploid hematopoietic stem cells and induced their differentiation *in vitro* using a cocktail of cytokines (Fig. 4A). We obtained a population of primary cells composed of diploid hematopoietic stem cells / progenitors and polyploid megakaryocytes showing different cell and nuclear sizes. These differences in size reflect a ploidy continuum and allow us to identify polyploid cells (Supp. Fig. 9A-B). Using live imaging approaches of megakaryocytes isolated from mice expressing H2B-mCherry, we observed that nuclei exit mitosis without chromosome segregation or following limited movement during anaphase A. These lack any sign of cytokinesis furrowing confirming MS (Fig. 4B and Movie 9-10), as previously suggested^7,60^.

**Figure 4:**
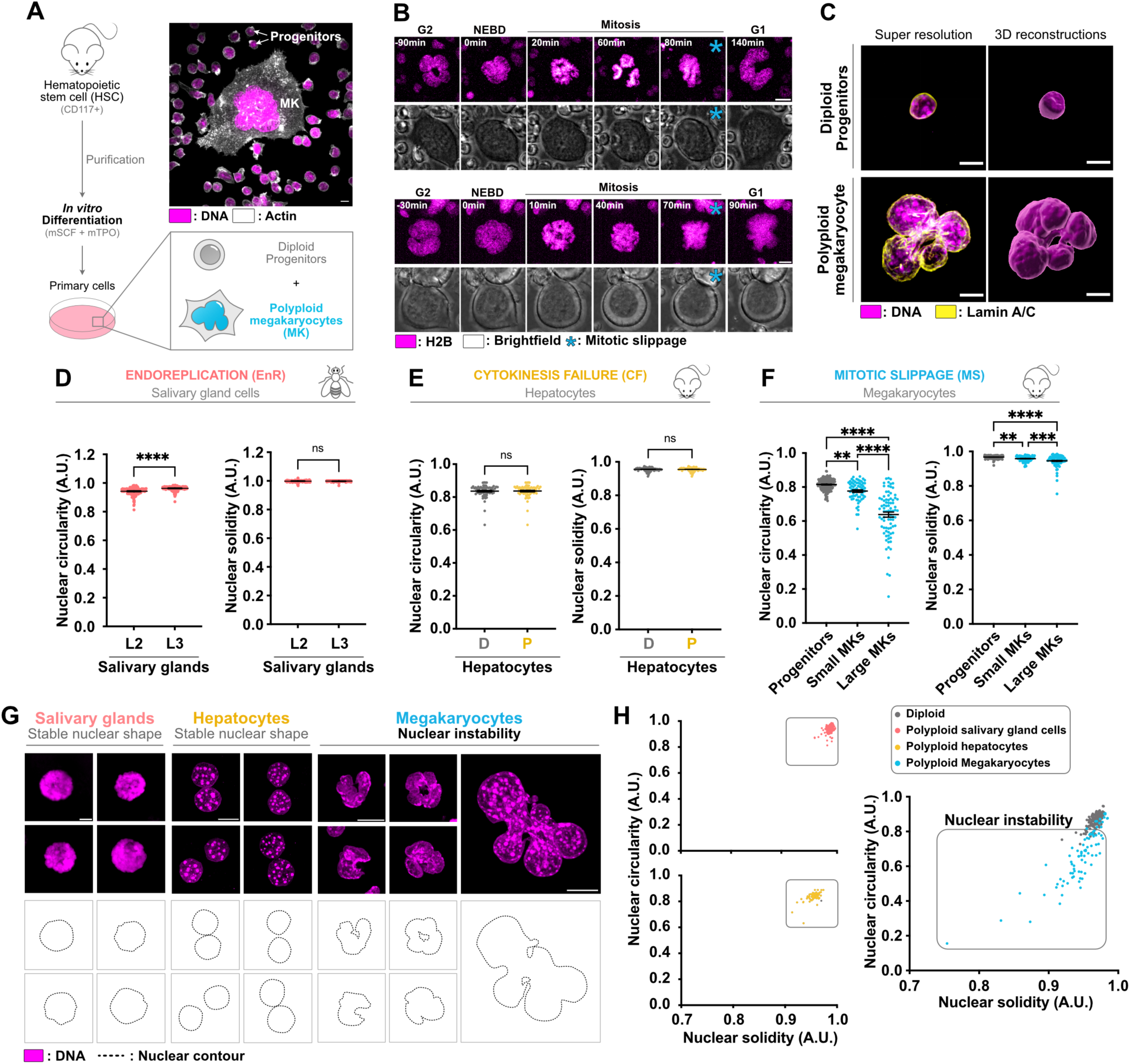
Megakaryocytes undergo mitotic slippage and display nuclear instability **(A)** Left panel - schematic workflow showing the methods used to induce megakaryocyte (MK) differentiation *in vitro*. Right panel – representative image showing a large polyploid megakaryocyte surrounded by diploid progenitors. Actin in grey, DNA in magenta. **(B)** Stills from time-lapse imaging of MKs stably expressing mCherry-H2B (in magenta). Brightfield in grey. **(C)** SIM representative images and corresponding 3D reconstructions showing diploid progenitors and polyploid megakaryocyte. Lamin A/C in yellow, DNA in magenta. **(D-F)** Graphs showing nuclear circularity and solidity in **(D)** *Drosophila* salivary glands (in red), in **(E)** mouse diploid (in grey) or polyploid hepatocytes (in yellow) and in **(F)** diploid mouse progenitors (in grey) and in polyploid megakaryocytes (in blue). Mean ± SEM, **(D)** >200, **(E)** >65, **(F)** >65 interphase cells analysed per condition, three independent experiments. **(G)** Representative images of megakaryocyte, hepatocyte or salivary gland nuclei, the nuclear contour is indicated below. DNA in magenta. **(H)** Graph showing the nuclear circularity and solidity index from **(D-F)** in *Drosophila* salivary glands (in red), in mouse diploid (in grey) or polyploid hepatocytes (in yellow) and in diploid mouse progenitors (in grey) and in polyploid megakaryocytes (in blue). Scale bars: 10µm. MS=mitotic slippage. EnR=endoreplication. CF=cytokinesis failure. MK=megakaryocytes. NEBD=nuclear envelope breakdown. A.U.=arbitrary unit. **(D-E)** t-test (two-sided). **(F)** ANOVA test (one-sided). ns=not significant. *=P ≤ 0.05. **= P ≤ 0.01. ***= P ≤ 0.001. ****= P ≤ 0.0001.

In our hands, and in agreement with *in vivo* analysis^6,7^, megakaryocytes displayed a single multilobulated highly deformed nucleus (Fig. 4C). Quantification of circularity and solidity revealed a decrease in a ploidy-dependent manner (Fig. 4D-F). Similarly to MS-induced RPE-1 cells, we observed that megakaryocyte nuclear shapes were highly heterogenous (Fig 4G-H). This reflects that megakaryocyte polyploidization by MS also generates nuclear instability. When comparing with other physiological polyploid cells, we noticed that nuclear instability was specific to megakaryocytes. Indeed, analysis of polyploid hepatocytes isolated from mouse liver, that become polyploid by CF^12,13^ or *Drosophila* salivary glands, that become polyploid by EnR^8,10^, showed homogeneous circular nuclei lacking nuclear deformations (Fig 4D-H and Supp. Fig. 9C-I). Similar observations were made *in vivo* in two other polyploid *Drosophila* tissues – the fat body and the ring gland that become polyploid by EnR^3^ (Supp. Fig. 9J).

Together, these findings show that MS, but not CF or EnR impact nuclear shape and stability in physiological polyploid settings.

### Mitotic slippage in megakaryocytes is associated with changes in chromatin compaction and nuclear envelope composition

Our results in RPE-1 cells show that MS-induced tetraploid cells exhibit changes in chromatin compaction and nuclear envelope composition promoting nuclear softness and deformability facilitating nuclear deformations by the microtubules. We therefore considered the possibility that megakaryocytes undergo a similar process.

We first tested whether histone compaction was altered upon MS in megakaryocytes. We found an increase in H3K27ac and H3K9ac levels in small megakaryocytes (Fig. 5A-B) while H3K14ac levels were increased in both small and large megakaryocytes (Supp. Fig. 10A). We also observed a global decrease in H3K9me2 levels with megakaryocyte ploidy while H3K9me3 levels remained comparable between progenitors and megakaryocytes (Supp. Fig. 10B-C). Interestingly, a local decrease in H3K9me2 levels in deformed regions was also found in megakaryocytes using two different antibodies (Supp. Fig. 10E-H). Concerning the nuclear envelope composition, a striking switch in Lamina composition during megakaryocyte differentiation was observed. Small and large megakaryocytes displayed lower levels of Lamin A/C than diploid progenitors, while Lamin B1 levels remained largely unchanged (Fig. 5C-D and Supp. Fig. 10D). In contrast, defects in H3K9me2 distribution and Lamina composition in hepatocytes or in the Lamina network in salivary gland cells were not detected (Supp. Fig. 10J-R). The high levels of histone acetylation and the decrease in Lamin A/C levels in megakaryocytes – two phenotypes associated with softer nuclei^43^ - are consistent with previous data showing that nuclear stiffness decreased during megakaryocyte differentiation^61^. To confirm that low nuclear stiffness contributes to nuclear deformations in megakaryocytes we promoted nuclear compaction by treating cells with Methylstat using a similar strategy as described above for RPE-1 cells. This was sufficient to alter nuclear shape and to reduce nuclear instability in megakaryocytes (Fig. 5E-G). Together, these findings suggest that during differentiation, megakaryocytes are in a state in which nuclear resistance is low - due to changes in chromatin compaction and nuclear envelope composition - favouring nuclear deformability. We next investigated whether the microtubule cytoskeleton also contribute to the nuclear deformations in megakaryocytes. We found thick microtubule bundles positioned around nuclear invaginations (Supp. Fig. 10I), which were absent in hepatocytes or salivary glands. In these former examples, the microtubule cytoskeleton was organized around the nucleus forming a microtubule ring (Supp. Fig. 10S). Importantly, removing microtubules was sufficient to decrease the nuclear instability in megakaryocytes (Supp. Fig. 10T), similarly to what was observed in RPE-1 MS-induced tetraploid cells. These results suggest that highly specialized and differentiated cells such as polyploid megakaryocytes rely on low nuclear stiffness to generate cells with irregular nuclear architecture *in vivo*. Importantly, our findings also identified the molecular mechanism underlying the irregular nuclear shape in megakaryocytes, initially described 50 years ago^6^.

**Figure 5:**
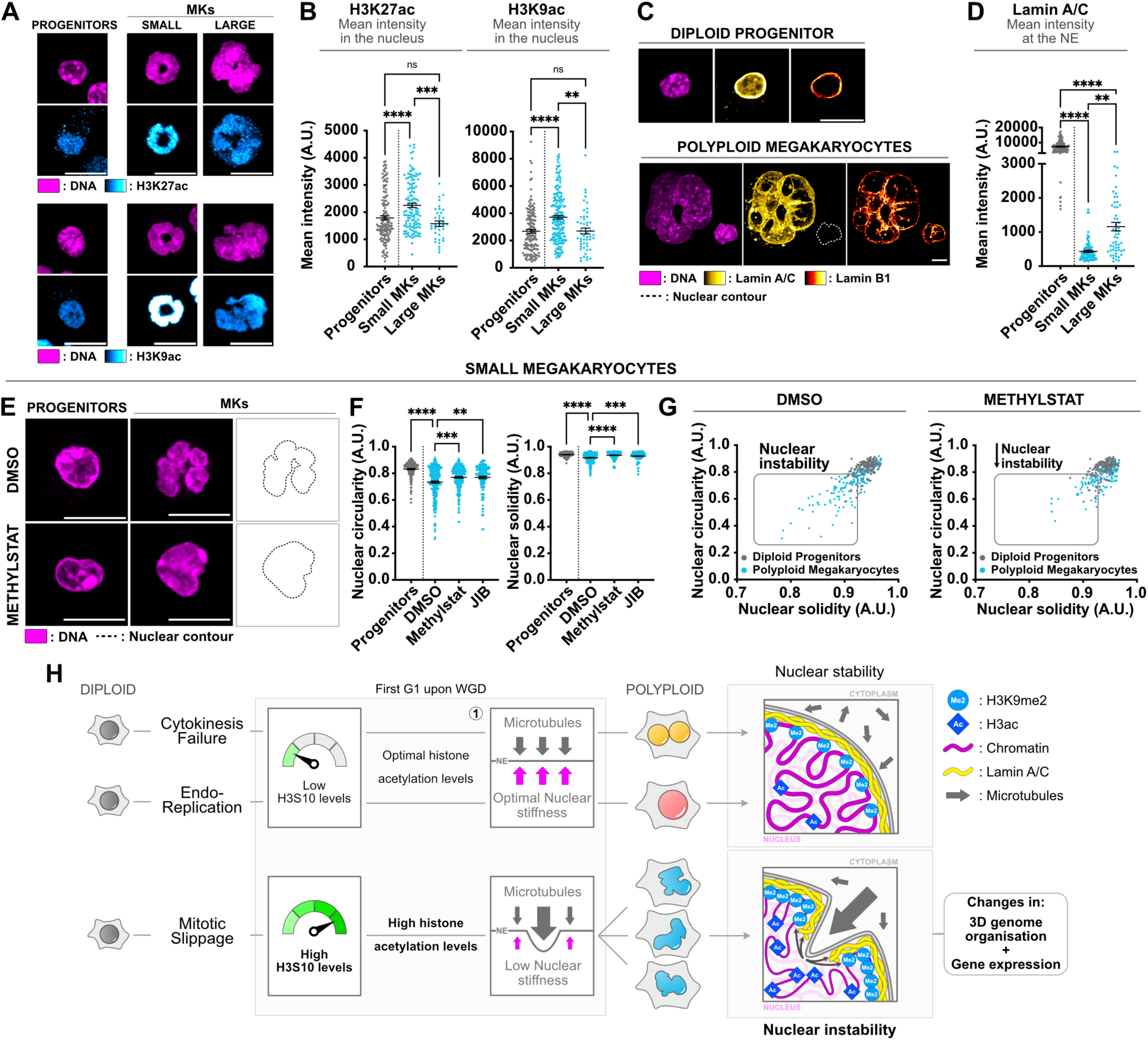
Mitotic slippage in megakaryocytes alters chromatin and nuclear envelope composition, promoting nuclear deformation **(A)** Representative images of mouse diploid progenitors and polyploid megakaryocytes. DNA in magenta. H3K27ac (upper panel) and H3K9ac (lower panel) in cyan hot. **(B)** Graph showing the H3K27ac (left panel) and H3K9ac (right panel) levels in diploid (in grey) and in polyploid megakaryocytes (in blue). Mean ± SEM, >140 interphase cells analysed per condition, three independent experiments. **(C)** Representative images of diploid progenitor and polyploid megakaryocytes. Lamin A/C in yellow hot, Lamin B1 in red hot, DNA in magenta. The dotted line shows the nuclear contour. **(D)** Lamin A/C mean intensity at the nuclear envelope in diploid (in grey) and in polyploid megakaryocytes (in blue). Mean ± SEM, >150 interphase cells analysed per condition, three independent experiments. **(E)** Representative images of diploid progenitors and polyploid megakaryocytes treated with DMSO or Methylstat. DNA in magenta. The dotted line shows the nuclear contour. **(F)** Graph showing the nuclear circularity and solidity indices in diploid progenitors (in grey) and polyploid megakaryocytes (in blue) treated with DMSO or Methylstat or JIB. Mean ± SEM, >90 interphase cells analysed per condition, three independent experiments. **(G)** Graph showing the nuclear circularity and solidity index from **(F)** in diploid progenitors (in grey) and polyploid megakaryocytes (in blue) treated with DMSO or Methylstat. **(H)** MS, unlike CF and EnR, generated tetraploid cells displaying abnormal nuclear configurations. These defects include maintenance of high H3S10 phosphorylation levels in G1, which promotes histone acetylation and thus decreased nuclear stiffness. Scale bars: 10µm. MK=megakaryocyte. NE=nuclear envelope. A.U.=arbitrary unit. **(B,D,F)** ANOVA test (one-sided). ns=not significant. *=P ≤ 0.05. **= P ≤ 0.01. ***= P ≤ 0.001. ****= P ≤ 0.0001.

## Discussion

In this study, we show that different routes to WGD lead to different outcomes. Indeed, MS, unlike CF and EnR, generated tetraploid cells displaying abnormal nuclear configurations. These defects include maintenance of high H3S10 phosphorylation levels in G1, which seem to promote histone acetylation and thus decreased nuclear stiffness. Due to the increased nuclear deformability, cytoplasmic microtubules generate nuclear invaginations that coincide with local changes in the chromatin / nuclear envelope composition. These findings are supported by other studies, which have shown that microtubules, Lamins and peripheral heterochromatin also contribute to nuclear morphology in diploid cells^42,62–64^. Together, these defects produce a heterogenous population of polyploid nuclei with various shapes - that we defined as nuclear instability - culminating with changes in 3D genome organisation as well as in gene expression (Fig. 5H).

Importantly, we observed these defects in the first G1 upon MS contributing to a better understanding of the early consequences of WGD. However, while the extent of the nuclear deformations is visually impressive, the effects on 3D genome organisation and on gene expression remain mild, even if significant, in the first G1 upon MS. This suggest that tetraploid cells can, at least in part, deal with these defects. Nevertheless, we cannot exclude that these early changes may be amplified after subsequent cell cycles.

MS, CF and EnR have been proposed to be at the origin of WGD in tumours^2,12,24^, but little is known about their proportion and respective role in cancer. Nuclear deformations were observed in tumours since the mid-nineteenth century and are even used for diagnostic in some situations^65^, but a potential link with WGD and MS was never established. It remains to be understood whether nuclear deformations and instability, typical of MS, contribute to the tumorigenic process. MS, CF and EnR are also involved in developmental programs generating physiological polyploid cells^7,8,13,66^. Whether these different strategies reflect a particular advantage or are just byproducts of selective pressure remains unknown.

Interestingly, to our knowledge, megakaryocytes are the only example of MS-generated physiological polyploid cells, making them quite unique. By showing that megakaryocytes display changes in chromatin / nuclear envelope composition that contributes to nuclear deformations, we exposed a molecular mechanism underlying the generation of megakaryocytes with irregular nuclear architecture. It has been shown that megakaryocyte reprogramming to induce polyploidy through EnR instead of MS contributes to prevent nuclear deformations^60^, confirming the predominant role of MS in generating megakaryocytes with irregular architecture. The global changes in chromatin state that we observed upon MS may contribute to alterations in gene expression sustaining the required transcriptomic changes described during megakaryocyte differentiation^67^.

Overall, our findings show that the cell cycle route chosen to generate polyploid cells has different outcomes. More importantly, our results highlight the fact that in physiological and pathological conditions, different methods to induce WGD have the potential to generate polyploid cells with specific features contributing to their inherent function.

## Acknowledgments

The authors acknowledge the Cell and Tissue Imaging (PICT-IBiSA) from Institut Curie, member of the national infrastructure France-BioImaging (https://ror.org/01y7vt929) supported by the French National Research Agency (ANR-24-INBS-0005 FBI BIOGEN) and V. Fraisier, C. Guedj. M. David, M. Cortes and A.S. Macé (PICT-IBiSA) for their continuous support on image acquisition and analysis. We thank L. Guyonnet, A. Chipont, A.G. Lafont and C. Guerrin from the Cytometry platform of Institut Curie. High-throughput sequencing was performed by the ICGex NGS platform of the Institut Curie supported by the grants ANR-10-EQPX-03 (Equipex) and ANR-10-INBS-09-08 (France Génomique Consortium) from the Agence Nationale de la Recherche (“Investissements d’Avenir” program), by the ITMO-Cancer Aviesan (Plan Cancer III) and by the SiRIC-Curie program (SiRIC Grant INCa-DGOS-465 and INCa-DGOS-Inserm_12554). Data management, quality control and primary analysis were performed by the Bioinformatics platform of the Institut Curie. We thank Genosplice (http://www.genosplice.com) and more specifically P. Delagrange and M. Caron for RNAseq and ATACseq analysis and for their valuable work. We thank A. Rodriguez-Romera and B. Psaila for sharing unpublished work. We thank A. Poleshko and JA Epstein for sharing the plasmids expressing H3WT and H3S10A. We thank A. Donada, J. Cossgrove and L. Perie, for teaching us the secrets of megakaryocyte differentiation and together with M. Piel and U. Kutay for helpful discussions and/or comments on the manuscript. GA’s team has been supported by the European Research Council (ERC-2015-ADG-694694 ChromADICT), the Ligue Nationale contre le Cancer (Equipe labellisée Ligue), France and Agence Nationale de la Recherche, France (ANR-11-LABX-0044_DEEP, ANR-10-IDEX-0001-02 PSL). N.W has been supported by the FRM. This work has been funded by InCA (www.e-cancer.fr) (2021-1- PREV-Bio grant), the government through the Agence Nationale de la Recherche from France 2030 (ANR-23-CHBS-0012), the Institut Curie and the CNRS.

## Author’s contribution

S.G. and R.B. conceived the project. S.G. wrote the manuscript. S.G. and M.B. did most of the experiments and data analysis presented here. S.A., F.SdC., C.V. and V.M. contributed to the HSC purification. A.S. prepared the RNA used for the RNAseq.

Y.M. and S.W. performed the AFM with the help of S.G. and M.B. J.F. and C.D. contributed by isolating the mouse hepatocytes. R.L., G.C. and E.O. performed the HiC experiments. N.W, A.F. and G.A. designed and performed the ATACseq experiments with the help of S.G. and M.B. R.B. obtained the funding used to develop this project. All authors read and commented on the manuscript.

## METHODS

### Human cell culture and generation of cell lines

#### Cell culture

Cells were maintained at 37°C in a 5% CO_2_ atmosphere. hTERT RPE-1 cells (female, ATCC Cat# CRL-4000, RRID:CVCL_4388), HEK293T cells (female, ATCC Cat# CRL-1573, RRID:CVCL_0045), BJ cells (male, ATCC Cat# CRL-4001, RRID:CVCL_6573) and HCT116 cells (male, ATCC Cat# CCL-247, RRID:CVCL_0291) were grown in Dulbecco’s modified medium (DMEM) F12 (11320-033 from Gibco) containing 10% fetal bovine serum (GE Healthcare), 100 U/ml penicillin, 100 U/ml streptomycin (15140-122 from Gibco).

All cells were routinely checked and tested negative for mycoplasma infection. Identity and purity of the human cell lines used in this study were tested and confirmed using STR authentication.

#### Generation of RPE-1 GFP-H3 WT and GFP-H3S10A stable cell lines

RPE-1 cells were transfected with 10µg GFP-H3 WT and GFP-H3S10A^1^ using JET PRIME kit (Polyplus Transfection, 114-07) according to the manufacturer’s protocol. After 24hrs, 500µg/ml G418 (4727878001 from Roche) was added to the cell culture medium. After 72hrs of antibiotic selection GFP-positive cells were then collected using Sony SH800 FACS (BD FACSDiva Software Version 8.0.1).

GFP-H3 WT and GFP-H3S10A plasmids were a gift of Epstein JA^1^.

#### Generation of RPE-1 GFP-Lamin and mCherry H2B stable cell line

RPE-1 mCherry cells^2^ were transfected with 10µg GFP Lamin chromobodies (Icg from Chromotek) using JET PRIME kit (Polyplus Transfection, 114-07) according to the manufacturer’s protocol. After 24hrs, 500µg/ml G418 was added to the cell culture medium. After 72hrs of antibiotic selection mCherry-H2B and GFP-positive (Lamin chromobodies) cells were then collected using Sony SH800 FACS (BD FACSDiva Software Version 8.0.1).

### Transgenic Animals

For animal care, we followed the European and French National Regulation for the Protection of Vertebrate Animals used for Experimental and other Scientific Purposes (Directive 2010/63; French Decree 2013-118). The project was authorized and benefited from guidance of the Animal Welfare Body, Research Centre, Curie Institute. All mice were kept in the Curie Institute Specific Pathogen Free (SPF) animal facility for breeding. The transgenic R26-H2B-mCherry mouse line^3^ was back-crossed in our animal facility on a C57Bl6/N genetic background (Charles River, France). Only heterozygous R26-H2B-mCherry +/- and control littermate males between 3 and 6 months of age were included in the analysis. Animals were anaesthetized by inhalation of isoflurane (Baxter, DDG9621) before being sacrificed by cervical dislocation for bone tissue collection.

### Mouse Megakaryocyte *in vitro* differentiation and cell culture

#### Hematopoietic progenitor enrichment

Tibias and femurs from 2 to 6-months old mice were dissected from the mouse and excess tissue was removed using scalpels. One epiphyse of each bone was cut ant the opened bone was placed into a 1.5mL centrifugation tube containing a 200uL tip. The bones were then centrifuged at high speed during 10 seconds in an Eppendorf tube to eject the bone marrow cells. The cells were resuspended in 1ml of cold RPMI containing 10% fetal bovine serum (GE Healthcare), 100 U/ml penicillin, 100 U/ml streptomycin (15140-122 from Gibco), Anti-Anti (11570486 from Gibco) - hereafter called complete RPMI medium - and centrifuged for 5min at 1700rpm, 4°C.

125µl of cold complete RPMI medium were used to resuspend the cells, then 25µl of anti-CD117 magnetic beads (130-097-146 from Miltenyi Biotec) were added and mix to the cells. Cells were incubated on ice for 30min with the beads. Then, 1ml of cold complete RPMI medium was added to the cells and they were filtered through a 100µm cell strainer (431752 from Coning) into 50ml tubes. The tubes and the cell strainers were washed with an additional 1ml of cold complete RPMI medium. Cells were centrifuged for 5min at 1700rpm, 4°C. During the centrifugation, a MS column (130-042-201 from Miltenyi Biotec) was prewet with 2ml of cold complete RPMI medium and installed on MACS magnet (130-042-501 from Miltenyi Biotec). The cells were then resuspended in 1ml of cold complete RPMI medium and transferred to a 2ml tube. Using an insulin syringe fitted with a short 25G needle (300400 from BD Bioscience), cells were transferred to the column. After, the tubes and the columns were washed using 2ml of cold complete RPMI medium. The CD117-positive fraction was then collected in a tube by pouring the cells using a plunger. The CD117-positive cells were centrifuge for 5min at 1700rpm, 4°C and resuspended in 500µl of cold complete RPMI medium.

#### Cell culture and In vitro differentiation

Purified CD117-positive cells were cultured on coverslips pre-treated with PolyLysine (0.02mg/ml during 5’, 37°C; HY-126437B from MedChemExpress) and Fibronectin (20µL in 500µl PBS, ON 37°C; F1141-5MG from Sigma-Aldrich) in DMEM-F12 + 10% fetal bovine serum (GE Healthcare) + 100 U/ml penicillin, 100 U/ml streptomycin (15140-122 from Gibco). Cells were maintained at 37°C in a 5% CO_2_ atmosphere. Megakaryocyte differentiation was induced by adding mSCF (50 ng/mL; 250-03 from Peprotech) + mTPO (50 ng/mL; 315-14 from Peprotech) for at least 3 days.

### Isolation of mouse Hepatocytes

Two old C57B6J mice were anesthetized with an intra-peritoneal injection of 10 mg/kg xylazine and 100 mg/kg ketamine. Primary hepatocytes were isolated by the two-step liver collagenase perfusion method and seeded in complete medium, as described previously^4^. Briefly, after caudal vena cava catheterization and portal vein opening, livers were washed with 50 mL of perfusion solution containing 10 mM 4-(2-hydroxyethyl)-1-piperazineethanesulfonic acid (HEPES, Merck #H6147), 137 mM NaCl (Merck #5886), 2.68 mM KCl (Merck #P5405), 232 µM Na2HPO4 (Merck #S5136), 0.5 mM ethylenediaminetetraacetic acid (EDTA, Thermo Fisher Scientific #15575020) dissolved in deionized sterile water. Then, livers were digested with a collagenase solution containing 0.5 mg/mL of collagenase type IV (from Clostridium histolyticum, Merck #C5138) diluted in perfusion solution, and 10 mM of CaCl2 (Merck #C7902). After isolation, hepatocytes were collected, washed, and resuspended in complete medium: William’s E medium GlutaMAX (Thermo Fisher Scientific #32551020) supplemented with bovine serum albumin 0,1% (Merck #A8412), penicillin-streptomycin 100 U/mL (Thermo Fisher Scientific #15140122), fungizone B 100 µg/mL (Thermo Fisher Scientific #15290026), dexamethasone 25 nM (Merck #D2915) and supplemented with 10% decomplemented foetal bovine serum (FBS, Thermo Fisher Scientific #10270106). The cell viability was equivalent in all groups (≥ 85%) as assessed by the Trypan blue exclusion method. Hepatocytes were seeded at a density of 6.0e+5 in 6-cm dishes and incubated at 37°C under an atmosphere containing 5% (v/v) CO2, for 4 hours, to allow the cells to adhere. The medium was then replaced with fresh FBS-free complete medium which remained unchanged for the 24h of primary culture.

### Fly husbandry and fly stocks

Flies were raised on cornmeal medium (0.75% agar, 3.5% organic wheat flour, 5.0% yeast, 5.5% sugar, 2.5% nipagin, 1.0% penicillin-streptomycin and 0.4% propionic acid). Fly stocks were maintained at 18 °C. Stocks used in this study: w[1118] (3605 from Bloomington).

In all experiments, larvae were staged to obtain comparable stages of development. Egg collection was performed at 25 °C for 24 h. After development at 25 °C, third instar larvae were used for dissection.

### Treatments

#### Chemicals

The drugs were used at the following concentrations: Auxin (I5148 from Sigma), 500µM; Doxycycline (D3447 from Sigma), 2µg/ml; Asunaprevir (S4935 from Selleckchem), 3µM; Monastrol (S8439 from Selleckchem), 50µM; MPI-0479605 (S7488 from Selleckchem), 1µM; Blebbistatin (B0560 from Sigma), 20µM; Palbociclib (S1579 from Selleckchem), 1µM; Methylstat (HY-15221 from Medchem), 2 (for live imaging or megakaryocytes) or 4µM (for fixed cells); JIB (HY-13953 from MedChemExpress), 2,5 (for megakaryocytes) or 5µM (for human cell lines), Nocodazole (HY-13520 from Medchem), 2µM; Taxol (T1912 from Sigma), 1nM; SP600125 (HY-12041 from Medchem), 20µM; RO3306 (HY-12529 from Medchem), 10µM; Calyculin A (HY-18983 from Medchem), 5nM; C646 (HY-13823 from Medchem), 30µM.

Methylstat, JIB and C646 were added 6hrs before the induction of whole genome duplication. Nocodazole and Taxol were added 2hrs before the induction of whole genome duplication.

Cells were treated with Calyculin A for 2:30hrs before fixation.

Megakaryocytes were incubated with Methylstat and JIB for 6hrs three days after the addition of the cytokines. To analyze megakaryocytes that performed mitotic slippage in the presence of the drugs only the small megakaryocytes were considered.

#### Induction of WGD in human cell lines

To induce mitotic slippage, cells were incubated with DMSO (D8418 from Sigma Aldrich) or with a combination of 50µM Monastrol (S8439 from Selleckchem) + 1µM MPI-0479605 (S7488 from Selleckchem) for at least 4hrs.

Alternatively, to induce mitotic slippage, CCNB1 depletion in RPE CCNB1^AID^ cells was induced as described before^5^. Briefly, cells were treated with 2µg/ml Doxycycline (D3447 from Sigma Aldrich) + 3µM Asunaprevir (S4935 from Selleckchem) for 2hrs. Then, 500 µM Auxin (I5148 from Sigma Aldrich) was added to the cell culture medium for at least 4hrs.

Mitotic slippage was induced using the combination of Monastrol + MPI-0479605 unless it is specified in the figure legends.

To induce cytokinesis failure, cells were incubated with DMSO (D8418 from Sigma Aldrich) or with 20µM Blebbistatin (B0560 from Sigma) or with 20µM Paprotrain (HY-101298 from Medchem) for at least 4hrs. Actin staining using Phaloidin (A30107 from ThermoFisher) was used to ascertain the binucleation of the CF-generated tetraploid cells.

Cytokinesis failure was induced using Blebbistatin unless it is specified in the figure legends.

To induce endoreplication, cells were incubated with DMSO (D8418 from Sigma Aldrich) or with 20µM SP600125 (HY-12041 from Medchem) for at least 4hrs.

Alternatively, to induce endoreplication, CCNA2 depletion in RPE CCNA2^AID^ cells was induced as described before^5^. Briefly, cells were treated with 2µg/ml Doxycycline (D3447 from Sigma Aldrich) + 3µM Asunaprevir (S4935 from Selleckchem) for 2hrs. Then, 500 µM Auxin (I5148 from Sigma Aldrich) was added to the cell culture medium for at least 4hrs.

Endoreplication was induced using SP600125 unless it is specified in the figure legends.

Each treatment generates a mixed population of diploid and tetraploid cells. After synchronizing cells in G1 (see the “Cell cycle synchronization” section), diploid (2N) and tetraploid (4N) cells are easily distinguished based on their nuclear size measured using Image J software V2.1.0/1.53c, https://imagej.net/software/fiji/downloads, as previously described^2^.

#### Cell cycle synchronization

To obtain a synchronous population of G1 cells, after 4hrs of WGD induction (see “Induction of WGD in human cell lines” section) 1µM Palbociclib (Cdk4/6 inhibitor, S1579 from Selleckchem) was added to the cell culture media for 16hrs to synchronize cells at the G1/S transition.

To analyze the first G2 upon WGD, Palbociclib was washed five times using PBS 1X and cells were released in the presence of 10µM RO-3306 (HY-12529 from Medchem) to block them in G2.

To analyze asynchronous cells, WGD induction was done during 4hrs before to fix the cells.

### Immunofluorescence

#### Preparation and imaging of human cells

Cells were plated on cover slips in 12-well plates and treated with the indicated drugs. To label cells, they were fixed using 4% of paraformaldehyde (15710 from Electron Microscopy Sciences) + Triton X-100 (2000-C from Euromedex) 0,1% in PBS (20 min at 4°C) or for alternative H3K9me2 staining with 100% cold Methanol (32213 from Honeywell) (10min at - 20°C). Then, cells were washed three times using PBS-T (PBS 1X + 0,1% Triton X-100 + 0,02% Sodium Azide) and incubated with PBS-T + BSA (04-100-812-C from Euromedex) 1% for 30 min at RT. After three washes with PBS-T + BSA, primary and secondary antibodies were incubated in PBS-T + BSA 1% for 1hr and 30 min at RT, respectively. After two washes with PBS, cells were incubated with 3 μg/ml DAPI (4’,6-diamidino-2-phenylindole; D8417 from Sigma Aldrich) for 15 min at RT. After two washes with PBS, slides were mounted using 1.25% n-propyl gallate (Sigma, P3130), 75% glycerol (bidistilled, 99.5%, VWR, 24388-295), 23.75% H2O.

Images were acquired on an upright widefield microscope (DM6B, Leica Systems, Germany) equipped with a motorized XY and a 40X objective (HCX PL APO 40X/1,40-0,70 Oil from Leica). Acquisitions were performed using Metamorph 7.10.1 software (Molecular Devices, USA) and a sCMOS camera (Flash 4V2, Hamamatsu, Japan). Stacks of conventional fluorescence images were collected automatically at a Z-distance of 0.5 µm (Metamorph 7.10.1 software; Molecular Devices, RRID:SCR_002368).

For H3.1/2 and H3.3 staining, two additional extractions were done, a pre-extraction (Triton X-100, 0,1%, 30sec) before the fixation and a post-extraction (Triton X-100, 0,5%, 5min) after the fixation.

#### Preparation and imaging of mouse megakaryocytes and mouse hepatocytes

For megakaryocytes, cells were plated on cover slips in 24-well plates and fixed using 4% of paraformaldehyde (15710 from Electron Microscopy Sciences) + Triton X-100 (2000-C from Euromedex) 0,1% in PBS (20 min at RT) or for microtubule staining with 100% cold MetOH (32213 from Honeywell) (10min-20°C). For hepatocytes, cells were plated on cover slips in 6-well plates and fixed using 4% of paraformaldehyde (15710 from Electron Microscopy Sciences) or for microtubule staining with 100% MetOH (32213 from Honeywell) (10min - 20°C).

Then, cells were washed three times using PBS-T (PBS 1X + 0,1% Triton X-100 + 0,02% Sodium Azide) and incubated with PBS-T + 1% BSA (04-100-812-C from Euromedex) 1% for 30 min at RT. After three washes with PBS-T + BSA, primary and secondary antibodies were incubated in PBS-T + BSA 1% for 3hrs and 2hrs at RT, respectively. After two washes with PBS, cells were incubated with 3 μg/ml DAPI (4’,6-diamidino-2-phenylindole; D8417 from Sigma Aldrich) for 15 min at RT. After two washes with PBS, slides were mounted using 1.25% n-propyl gallate (Sigma, P3130), 75% glycerol (bidistilled, 99.5%, VWR, 24388-295), 23.75% H2O.

Images were acquired on a spinning disk microscope (Gataca Systems, France). Based on a CSU-W1 (Yokogawa, Japan), the spinning head was mounted on an inverted Eclipse Ti2 microscope equipped with a motorized XY Stage (Nikon, Japan). Images were acquired through a 100x 1.4NA Plan-Apo objective with a sCMOS camera (Prime95B, Photometrics, USA). Optical sectioning was achieved using a piezo stage (Nano-z series, Mad City Lab, USA). Gataca Systems’ laser bench was equipped with 405, 491 and 561 nm laser diodes, delivering 150 mW each, coupled to the spinning disk head through a single mode fibre. Multi-dimensional acquisitions were performed using Metamorph 7.10.1 software (Molecular Devices, USA).

#### Preparation and imaging of Drosophila salivary glands

*Drosophila* salivary glands from third instar larvae were dissected in PBS and fixed for 30 min in 4% paraformaldehyde in PBS. They were washed 3 times in PBST 0.3% (PBS, 0.3% Triton X-100, T9284 from Sigma), 15 min for each wash, and incubated for 1 hr with blocking solution (PBS, 0.3% Triton X-100 + 5% normal goat serum, 10098792 from Gibco). Then salivary glands were washed 3 times in PBS, 15 min for each wash and incubated overnight at 4 °C with primary antibodies at the appropriate dilution in the blocking solution. Tissues were washed three times in PBST 0.3%, 15 min for each wash, and incubated overnight at 4 °C with secondary antibodies diluted in the blocking solution. Salivary glands were then washed 3 times in PBST 0.3%, 15 min for each wash, rinsed in PBS and incubated with 3 μg ml−1 DAPI (4′,6-diamidino-2-phenylindole; Sigma Aldrich D8417) at room temperature for 30 min. Salivary glands were then washed in PBS at room temperature for 30 min and mounted in clearing solution (Rapiclear 1.49 Sunjin Lab, RC149002).

Images were acquired as previously described in the section “*Preparation and imaging of mouse megakaryocytes and mouse hepatocytes”*.

#### Structural illumination microscopy

Images were acquired as previously described in the section “*Preparation and imaging of mouse megakaryocytes and mouse hepatocytes”*.

Super resolution was achieved on the CSU-W1 spinning disk equipped with a super-resolution module (Live-SR, Gataca systems). This module is based on structured illumination with optical reassignment technique and online processing leading to a two-time resolution improvement. The method called multifocal SIM (MSIM) allows combining the doubling resolution together with the physical optical sectioning of confocal microscopy.

#### Primary and secondary antibodies

Primary and secondary antibodies were used at the following concentrations: β-catenin (1/250; C2206 from Sigma-Aldrich, RRID:AB_476831), Lamin A/C (1/500; SAB4200236 from Sigma Aldrich, RRID:AB_10743057), Lamin B1 (1/500; ab16048 from Abcam, RRID:AB_443298), H3S10 (1/500, ab14955 from Abcam, RRID:AB_443110), Lamin A/C (1:50, ADL101 from DHSB, RRID:AB_528332), Lamin B (1/50, ADL67.10 from DHSB, RRID:AB_528336), H3K9ac (1/500, 9649S from Ozyme; RRID:AB_823528), H3K14ac (1/500, ab52946 from Abcam, RRID:AB_880442); H3K27ac (1/500, 39685 from Active Motif, RRID:AB_2793305), DM1A (1/1000, T9026 from Sigma Aldrich, RRID:AB_477593), H3K9me2 (1/500, 39239 from Active Motif, RRID:AB_2793199), H3K9me2 (1/500, ab1220 from Abcam; RRID:AB_449854), H3K9me3 (1/500, ab8898 from Abcam, RRID:AB_306848), H3K27me3 (1/500, 39155 from Active motif, RRID:AB_2561020), HP1⍺ (1/500, ab109028 from Abcam, RRID:AB_10858495), Lap2β (1/250, 611000 from BD Bioscience, RRID:AB_398313), LBR (1/500, ab232731 from Abcam, RRID:AB_3246426), Nuclear Pore Complex (1/250, ab24609 from Abcam, RRID:AB_448181), Emerin (1/150, 10351-1-AP from ProteinTech, RRID:AB_2100056), goat anti-Rabbit IgG (H+L) Highly Cross-Adsorbed Secondary Antibody, Alexa Fluor 647 (1/250; A21245 from ThermoFisher, RRID:AB_2535813), Goat anti-Rabbit IgG (H+L) Highly Cross-Adsorbed Secondary Antibody, Alexa Fluor 546 (1/250; A-11035 from ThermoFisher, RRID:AB_2534093), Goat anti-Mouse IgG1 Cross-Adsorbed Secondary Antibody, Alexa Fluor™ 488 (1/250, A-21121 from ThermoFisher, RRID:AB_2535764), Goat anti-Mouse IgG2b Cross-Adsorbed Secondary Antibody, Alexa Fluor™ 488 (1/250, A-21141 from ThermoFisher, RRID:AB_2535778), Goat anti-Mouse IgG2a Cross-Adsorbed Secondary Antibody, Alexa Fluor™ 546 (1/250, A-21133 from ThermoFisher, RRID:AB_2535772), Alexa Fluor™ Plus 647 Phalloidin (1/2000, A30107 from ThermoFisher).

### 3D cultures

#### Mitotic slippage on 3D cultures

To generate spheroids, 500 cells per well were seeded into 96 ultra-low attachment well plates (7007 from Corning) in presence of DMSO (D8418 from Sigma Aldrich) or with 50µM monastrol (S8439 from Selleckchem) and 1µM MPI-0479605 (S7488 from Selleckchem). Plates were spun down at 200g for 3min, to allow spheroid formation, and incubated for 24hrs at 37°C.

#### Immunostaining

Spheroids were collected and washed quickly with PBS prior fixation using 4% of paraformaldehyde (15710 from Electron Microscopy Sciences) in PBS for 40min. Then, spheroids were permeabilized for 5 min using Triton X-100 (2000-C from Euromedex) 0,3% in PBS and blocked for 30min using Blocking Buffer (BB= PBS 1X + 0,3% Triton X-100 + 0,02% Sodium Azide + 3% BSA). Aggregates were incubated with primary antibodies diluted into BB O/N. After 3 washes using BB, spheroids were incubated with secondary antibodies in BB for 3hrs. Cells were then washed several times for 2hrs in BB and mounted on glass with EverBrite (Biotium). For primary and secondary antibodies see *Immunofluorescence microscopy and antibodies* section.

### Time lapse microscopy

Cells were plated on a dish (627870 from Greiner). Methylstat (2µM) or Nocodazole (2µM) were added two hours before to start the movies. To follow microtubules or actin, SPY650-Tubulin (SC503 from Tebubio) or SPY650-FastAct (SC505 from Tebubio), respectively, were added one hour before to start the movie.

Images were acquired on a spinning disk microscope (Gataca Systems, France). Based on a CSU-W1 (Yokogawa, Japan), the spinning head was mounted on an inverted Eclipse Ti2 microscope equipped with a motorized XY Stage (Nikon, Japan). Images were acquired through a 40X NA 1.3 oil objective with a sCMOS camera (Prime95B, Photometrics, USA).

Optical sectioning was achieved using a piezo stage (Nano-z series, Mad City Lab, USA). Gataca Systems’ laser bench was equipped with 405, 491 and 561 nm laser diodes, delivering 150 mW each, coupled to the spinning disk head through a single mode fiber. Laser power was chosen to obtain the best ratio of signal/background while avoiding phototoxicity. Multi-dimensional acquisitions were performed using Metamorph 7.10.1 software (Molecular Devices, USA). Stacks of conventional fluorescence images were collected automatically at a Z-distance of 0.5 µm (Metamorph 7.10.1 software; Molecular Devices, RRID:SCR_002368). Images are presented as maximum intensity projections generated with ImageJ software (RRID:SCR_002285), from stacks deconvolved with an extension of Metamorph 7.10.1 software.

### FACS of diploid and tetraploid cells

A mix of diploid and tetraploid cells synchronized in G1 or in G2/M (see “Cell cycle synchronization” section) were incubated with 2 µg ml−1 Hoescht 33342 (Sigma Aldrich 94403) for 1 h at 37 °C, 5% CO2. Then, a single cell suspension was generated. Cells were washed using PBS, the supernatant was removed, and cells were resuspended in cell culture medium containing 1µM EDTA (10135423 from Invitrogen). Fluorescence-activated cell sorting (FACS) was performed using Sony SH800 FACS (BD FACSDiva Software Version 8.0.1), as previously described^2^. Compensations were performed using the appropriate negative control samples.

### HiC experiment

Diploid and tetraploid cells were isolated using flow cytometry (see “FACS of diploid and tetraploid cells” section). Diploid and tetraploid cells were isolated using flow cytometry (see “FACS of diploid and tetraploid cells” section). Library preparation for Hi-C was conducted as previously described^6^. Briefly, the cells were fixed with formaldehyde, and fixed nuclei were extracted for further processing. Chromatin was digested with MboI restriction enzyme (New England Biolabs, R0147), and digested ends were biotinylated. Next, a proximity ligation was carried out, followed by chromatin crosslink reversal. DNA was purified, sheared via sonication, and size-selected for library preparation. Biotin-marked DNA fragments were pulled down with Dynabeads MyOne Streptavidin C1 (Thermo Fisher Scientific, 65001). Hi-C library preparation continued with end polishing, A-tailing, and Illumina TruSeq unique dual indexes ligation reactions. Lastly, libraries were amplified via PCR, and sequenced on an Illumina HiSeq X platform, PE150 configuration.

### Atomic force microscopy

Atomic Force Microscopy measurements were performed on glass bottom dishes (ibidi) with a JPK NanoWizard 4 (Bruker Nano) microscope mounted on an Eclipse Ti2 inverted fluorescent microscope (Nikon) and operated via JPK SPM Control Software v.6. MLCT triangular silicon nitride cantilevers (Bruker) were used to access the mechanical properties of nuclei. Forces of up to 2 nN were applied at 10 micron per second constant cantilever velocity also ensuring an indentation depth of at ∼500 nm.

### ATACseq

#### Sample preparation

The samples were prepared using the following kit 53150 from Active Motif and according to the manufacturer’s protocol. We isolated diploid and tetraploid G1 cells using flow cytometry (see “FACS of diploid and tetraploid cells” section) and used 1 x10^5^ cells per condition. Spike in (53154 from Active Motif) was added according to the manufacturer’s protocol.

#### Sequencing

The pool was quantified by qPCR using the KAPA library quantification kit (Roche). Sequencing was carried out on the NovaSeq X Plus instrument from Illumina based on a 2*100 cycle mode (paired-end reads, 100 bases) targeting around 50 million clusters per sample.

### Total RNAseq

#### RNA preparation

Diploid and tetraploid G1 cells were isolated using flow cytometry (see “FACS of diploid and tetraploid cells” section). Then RNA was extracted using All Prep DNA/RNA Mini Kit (80204 from Qiagen) according to the manufacturer’s protocol. To normalize, RNA Spike-in were added to the samples (4456740 from Invitrogen) according to the manufacturer’s protocol. 1 x 10^6^ cells were used per condition (3n).

To control the quality of the samples before High-Throughput Sequencing, RNAs were analyzed to quantify the RIN using Agilent RNA 6000 Nano Kit (5067-1511 from Agilent).

#### Sequencing

We prepared RNA sequencing libraries from 500ng of total RNA using the Illumina® Stranded Total RNA Prep and the Ligation with Ribo-Zero Plus library preparation kit, which allows to perform a strand specific sequencing. This protocol includes a first step of enzymatic depletion of abundant transcripts from multiple species using specific probes. We then perfomed cDNA synthesis and used resulting fragments for dA-tailing followed by ligation of RNA Index Anchors. PCR amplification with indexed primers (IDT for Illumina RNA UD Indexes) was finally achieved, with 13 cycles, to generate the final cDNA libraries. Individual library quantification and quality assessment was performed using Qubit fluorometric assay (Invitrogen) with dsDNA HS (High Sensitivity) Assay Kit and LabChip GX Touch using a High Sensitivity DNA chip (Perkin Elmer). Libraries were then equimolarly pooled and quantified by qPCR using the KAPA library quantification kit (Roche). Sequencing was carried out on the NovaSeq X Plus instrument from Illumina using paired-end 2 x 100 bp, to obtain around 100 million clusters (200 million raw paired-end reads) per sample.

## QUANTIFICATION AND STATISTICAL ANALYSIS

### Immunofluorescence

Image analysis and quantifications were performed using Image J software V2.1.0/1.53c, https://imagej.net/software/fiji/downloads^7^. Images are presented as maximum intensity projections generated with ImageJ software (RRID:SCR_002285).

For the 3D reconstruction, images were converted in Imaris files using ImarisFileConverter 9.9.1 (Bitplane, RRID:SCR_007370). Then, images were imported into Imaris software v.9.6.0 (Bitplane, RRID:SCR_007370). Nuclear shapes were reconstructed based on the DNA staining. 3D views were generated using Imaris software and saved as TIFF.

To quantitatively measure nuclear shapes, the “wand” tool from Image J V2.1.0/1.53c software was used to select the nuclei based on the DAPI staining and then the circularity and solidity indexes were measured using the “shape descriptors” tool from Image J V2.1.0/1.53c software. To quantify fluorescence intensity, images were converted into maximum intensity projections and the mean fluorescence intensity at the whole nucleus was measured using the “polygon sections” tool from Image J V2.1.0/1.53c software while the mean fluorescence intensity at the nuclear envelope or locally at the intact/deformed regions were measured using the “segmented lines” tool from Image J V2.1.0/1.53c software. Since diploid cells were not deformed, the mean fluorescence intensity was compared at two random regions to visualize the distribution of the signal, while in newly born tetraploid cells, intact and deformed areas were compared.

The coefficient of variation (CoV) was measured by dividing the standard deviation of the signal by the mean fluorescence intensity for each cell.

To define small and large megakaryocytes or hepatocytes, the median nuclear size was used as a threshold.

For the figures, images were processed on Image J V2.1.0/1.53c software, and mounted using Affinity Designer 2, https://affinity.serif.com/fr/designer/.

### Time lapse microscopy

Image analysis and quantifications were performed using Image J software V2.1.0/1.53c, https://imagej.net/software/fiji/downloads. Videos were corrected using “3D correct drift” plugin with Image J V2.1.0/1.53c software to keep the cell of interest at the center of the region of interest.

For the Figure 2D, nuclei masks were generated using the Fiji Plugin as described in^8^. Briefly, this process involves denoising (using the Pure Denoise Fiji Plugin^9^) and thresholding the image over time, to get one ROI (the nucleus) on each frame. A custom macro was then used to project the contours over time, with each contour colored according to its corresponding frame. The Analyze Particles function was applied to the masks generated by the Plugin, and the contours’ ROIs were color-coded according to their frame using BAR Plugin.

To analyze nuclear shapes by live imaging, single cells were followed, and nuclear circularity and solidity indexes were measured in G2 before the NEBD and in G1 after complete DNA decondensation. The results were plotted using the “before-after” representation from GraphPad Prism (RRID:SCR_002798) version 10 for Mac, GraphPad Software, La Jolla California USA, www.graphpad.com. The delta circularity was measured by dividing the nuclear circularity index in G2 by the one in G1 for each single cell.

For the figures, images were processed on Image J V2.1.0/1.53c software, and mounted using Affinity Designer 2, https://affinity.serif.com/fr/designer/.

### Total RNAseq data

Analysis of sequencing data quality, reads repartition (e.g., for potential ribosomal contamination), inner distance size estimation, genebody coverage, strand-specificity of library were performed using FastQC, Picard-Tools, Samtools, and RSeQC. Reads were mapped using STAR on the Human hg38 genome assembly and read count was performed using featureCount from SubRead and the Human FAST DB v2022_1 annotations.

Gene expression was estimated as described previously^10^. Only genes expressed in at least one of the two compared conditions were analyzed further. Genes were considered as expressed if their FPKM value was greater than FPKM of 98% of the intergenic regions (background). Analysis at the gene level was performed using DESeq2^11^. Genes were considered differentially expressed for fold-changes ≥ 1.5 and p-values ≤ 0.05.

Overrepresented analyses and GSEA^12^ were performed using WebGestalt^13^ merging results from up-regulated and down-regulated genes only, as well as all regulated genes. Pathways and networks were considered significant with p-values ≤ 0.05.

VolcanoPlots were created using ggplot2. In blue are genes that are differentially under-expressed in the Tetraploid condition, and in red those that are over-expressed (based on thresholds indicated after the use of DESeq2).

### ATACseq

Quality control was performed using FastQC and MultiQC. Sequencing reads were aligned to the human reference genome (hg38) using the BWA-MEM2 algorithm. Duplicate reads were marked using the Picard MarkDuplicates tool. Only reads with a mapping quality score ≥40 were retained, using Samtools. Spike-in normalization was conducted following the guidelines provided by ActiveMotif. Briefly, the number of reads uniquely mapped to the Drosophila (dm6) genome was quantified per sample. A normalization ratio was then calculated based on the lowest number of Drosophila-mapped reads among all samples. This ratio was subsequently applied to down sample the uniquely mapped reads to the human genome in each sample with Samtools. Samtools was also used to shift BAM files. Peak calling was performed with MACS3 callpeak. Peaks overlapping ENCODE blacklist regions were removed using subtractBed. Peaks with log2 fold change (log2FC) < 3 and-log10(FDR) < 2 were filtered out. Differential peak analysis was carried out using DiffBind in a replicate mode. Finally, peak annotation was performed using HOMER.

### Atomic force microscopy

Analyses were performed with JPK Data Processing Software v.6 (Bruker Nano) by 1) removing the offset from the baseline of raw force curves, 2) identifying the contact point and subtracting cantilever bending then 3) fitting the Hertz model with correct tip geometry to quantitate the reported Young’s Modulus of nuclei.

### HiC data

For Hi-C data, reads were mapped to the human hg19 reference genome. Generation of Hi-C contact maps and further analyses, including inter-chromosomal interactions, intercompartmental interactions, and conservation of insulating boundaries were performed as previously described^6^.

### Statistical analysis

The statistical significance of differences was calculated using GraphPad Prism (RRID:SCR_002798) version 10 for Mac, GraphPad Software, La Jolla California USA, www.graphpad.com. The statistical test used for each experiment is indicated in the figure legends. Each representative image originates from the analyzed dataset and are representative of the associated quantifications. ns=not significant. *=P ≤ 0.05. **= P ≤ 0.01. ***= P ≤ 0.001. ****= P ≤ 0.0001.

## SUPPLEMENTARY FIGURE LEGENDS

**Figure S1:**
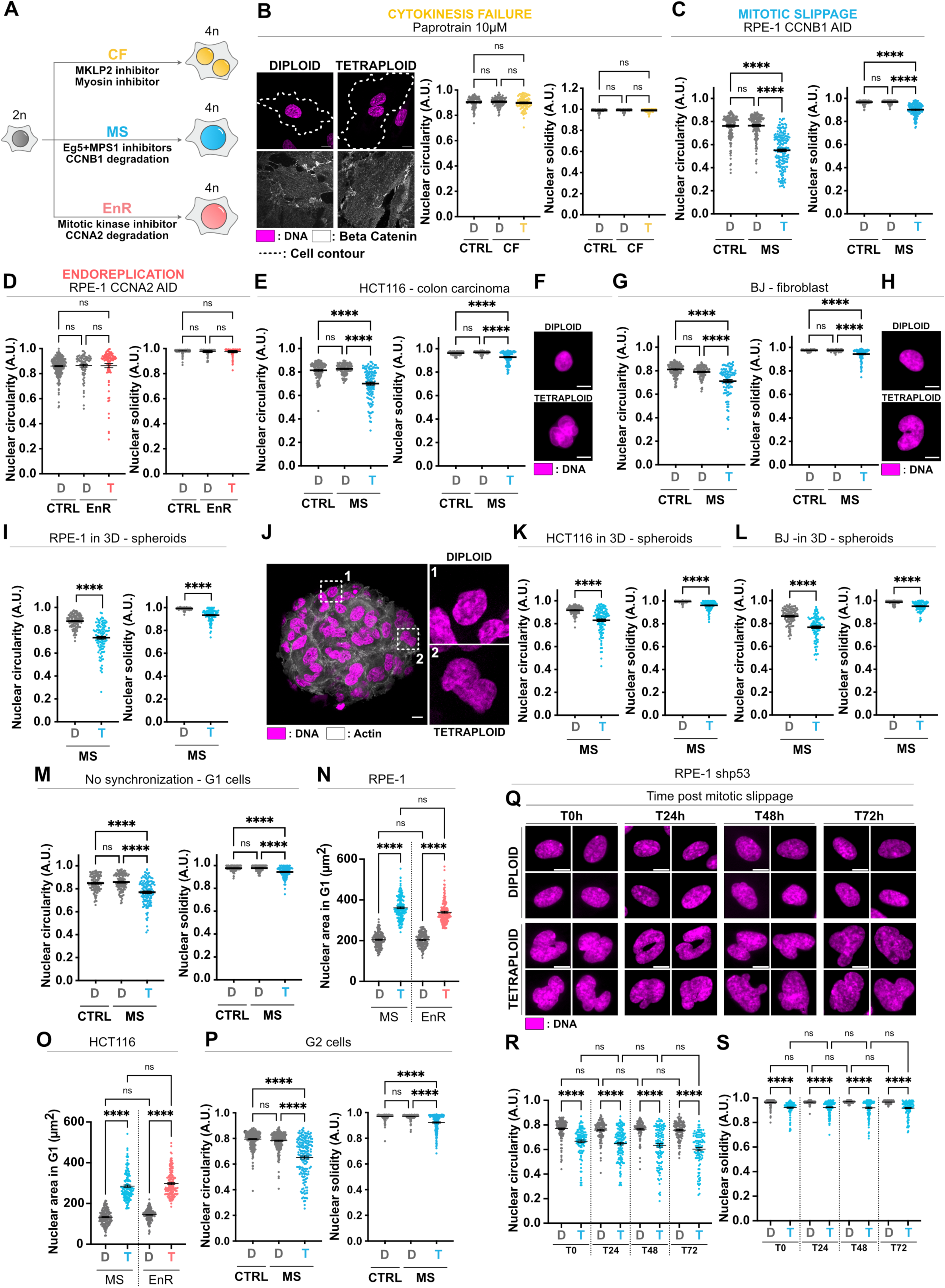
Mitotic slippage unlike cytokinesis failure or endoreplication, generates polyploid cells with abnormal nuclear architecture. **(A)** Schematic representation showing the methods used to induce whole genome duplication. **(B)** Left panel – representative images showing diploid and CF-generated tetraploid G1 RPE-1 cells. β-Catenin in grey, DNA in magenta. Right panel – Graph showing nuclear circularity and solidity in diploid and in CF-generated tetraploid G1 RPE-1 cells (in grey and yellow, respectively). Mean ± SEM, >100 G1 cells analysed per condition, three independent experiments. **(C-D)** Graphs showing nuclear circularity and solidity in diploid and in **(C)** MS- and **(D)** EnR-generated tetraploid G1 RPE-1 cells (in blue and red, respectively). Mean ± SEM, >150 G1 cells analysed per condition, three independent experiments. **(E)** Graphs presenting nuclear circularity and solidity in diploid and in MS-generated tetraploid HCT116 cells in G1 (in grey and blue, respectively). Mean ± SEM, >100 G1 cells analysed per condition, three independent experiments. **(F)** Representative images showing diploid and MS-generated tetraploid HCT116 nuclei. DNA in magenta. **(G)** Graphs showing nuclear circularity and solidity in diploid and in MS-generated tetraploid BJ cells in G1 (in grey and blue, respectively). Mean ± SEM, >100 G1 cells analysed per condition, three independent experiments. **(H)** Representative images presenting diploid and MS-generated tetraploid BJ nuclei. DNA in magenta. **(I)** Graph showing nuclear circularity and solidity in diploid and in MS-generated tetraploid RPE-1 cells cultured in 3D (in grey and blue, respectively). Mean ± SEM, >100 G1 cells analysed per condition, three independent experiments. **(J)** Representative images showing an RPE-1 spheroid containing diploid and MS-generated tetraploid cells. Actin in grey, DNA in magenta. **(K-L)** Graphs showing nuclear circularity and solidity in diploid and in MS-generated tetraploid **(K)** HCT116 and **(L)** BJ cells cultured in 3D (in grey and blue, respectively). Mean ± SEM, >110 G1 cells analysed per condition, three independent experiments. **(M)** Graph showing nuclear circularity and solidity in diploid and in MS-generated tetraploid asynchronous RPE-1 cells (in grey and blue, respectively). Mean ± SEM, >110 interphase cells analysed per condition, three independent experiments. **(N-O)** Graphs presenting nuclear area in diploid and in MS- and EnR-generated tetraploid **(N)** G1 RPE-1 and **(O)** HCT116 cells (in grey, blue and red, respectively). Mean ± SEM, >150 G1 cells analysed per condition, three independent experiments. **(P)** Graph showing nuclear circularity and solidity in diploid and in MS-generated tetraploid G2 RPE-1 cells (in grey and blue, respectively). Mean ± SEM, >170 G2 cells analysed per condition, three independent experiments. **(Q)** Representative images showing diploid and MS-generated tetraploid RPE-1 shP53 cells. DNA in magenta. **(R-S)** Graphs showing nuclear **(R)** circularity and **(S)** solidity in diploid and in MS-generated tetraploid RPE-1 shp53 cells (in grey and blue, respectively). Mean ± SEM, >100 interphase cells analysed per condition, three independent experiments. Scale bars: 10µm. D=diploid. T=tetraploid. MS=mitotic slippage. CF=cytokinesis failure. EnR= endoreplication. A.U.=arbitrary unit. **(B,C,D,E,G,M,N,O,P,R,S)** Anova-test (one sided). **(I,K,L)** t-test (two-sided). ns=not significant. *=P ≤ 0.05. **= P ≤ 0.01. ***= P ≤ 0.001. ****= P ≤ 0.0001.

**Figure S2:**
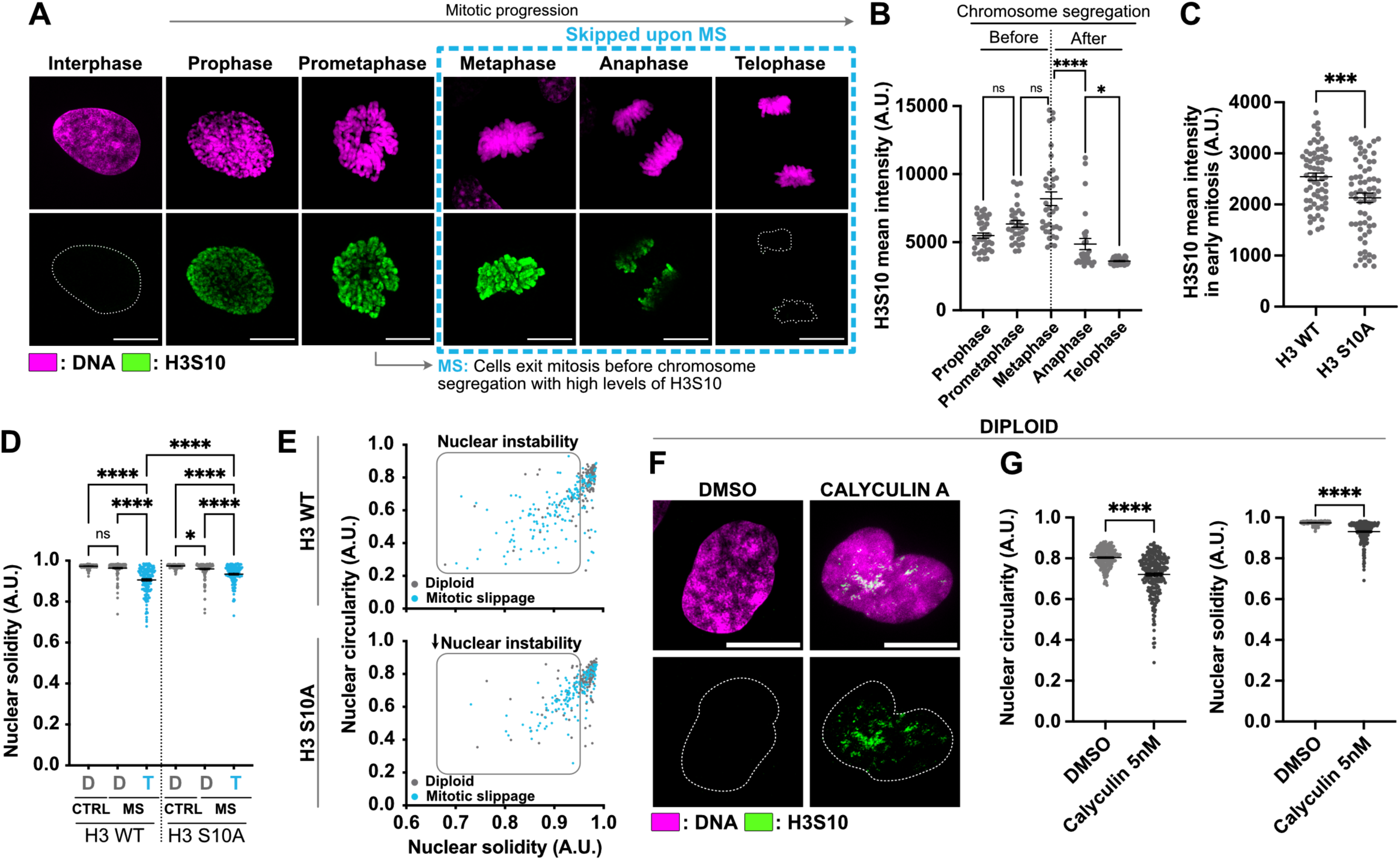
High levels of H3S10 phosphorylation contributes to nuclear deformations upon mitotic slippage. **(A)** SIM representative images showing diploid RPE-1 mitotic cells. H3S10 in green, DNA in magenta. **(B)** Graph showing H3S10 mean intensity during mitosis in diploid RPE-1 cells. Mean ± SEM, >25 mitotic cells analysed per condition, three independent experiments. **(C)** Graph showing the H3S10 mean intensity in RPE-1 diploid mitotic cells stably expressing H3 WT and H3 S10A. Mean ± SEM, >65 mitotic cells analysed per condition, three independent experiments. **(D)** Graph showing the nuclear solidity in diploid (in grey) and MS-generated tetraploid (in blue) G1 RPE-1 cells stably expressing H3 WT or H3 S10A. Mean ± SEM, >150 cells analysed per condition, three independent experiments. **(E)** Graphs showing the nuclear circularity and solidity index from **(D and Fig. 2D)** in diploid (in grey) and in tetraploid G1 RPE-1 cells (in blue) generated by MS. **(F)** SIM representative images showing diploid cells treated with DMSO or 5nM Calyculin. H3S10 in green, DNA in magenta. **(G)** Graphs showing the nuclear circularity and solidity index in diploid RPE-1 cells treated with DMSO or 5nM Calyculin. Mean ± SEM, >230 G1 cells analysed per condition, three independent experiments. Scale bar: 10µm. D=diploid. T=tetraploid. MS=mitotic slippage. NE= nuclear envelope. A.U.=arbitrary unit. (**B,D**) ANOVA test (one-sided). (**C,G**) t-test (two-sided). ns=not significant. *=P ≤ 0.05. **= P ≤ 0.01. ***= P ≤ 0.001. ****= P ≤ 0.0001.

**Figure S3:**
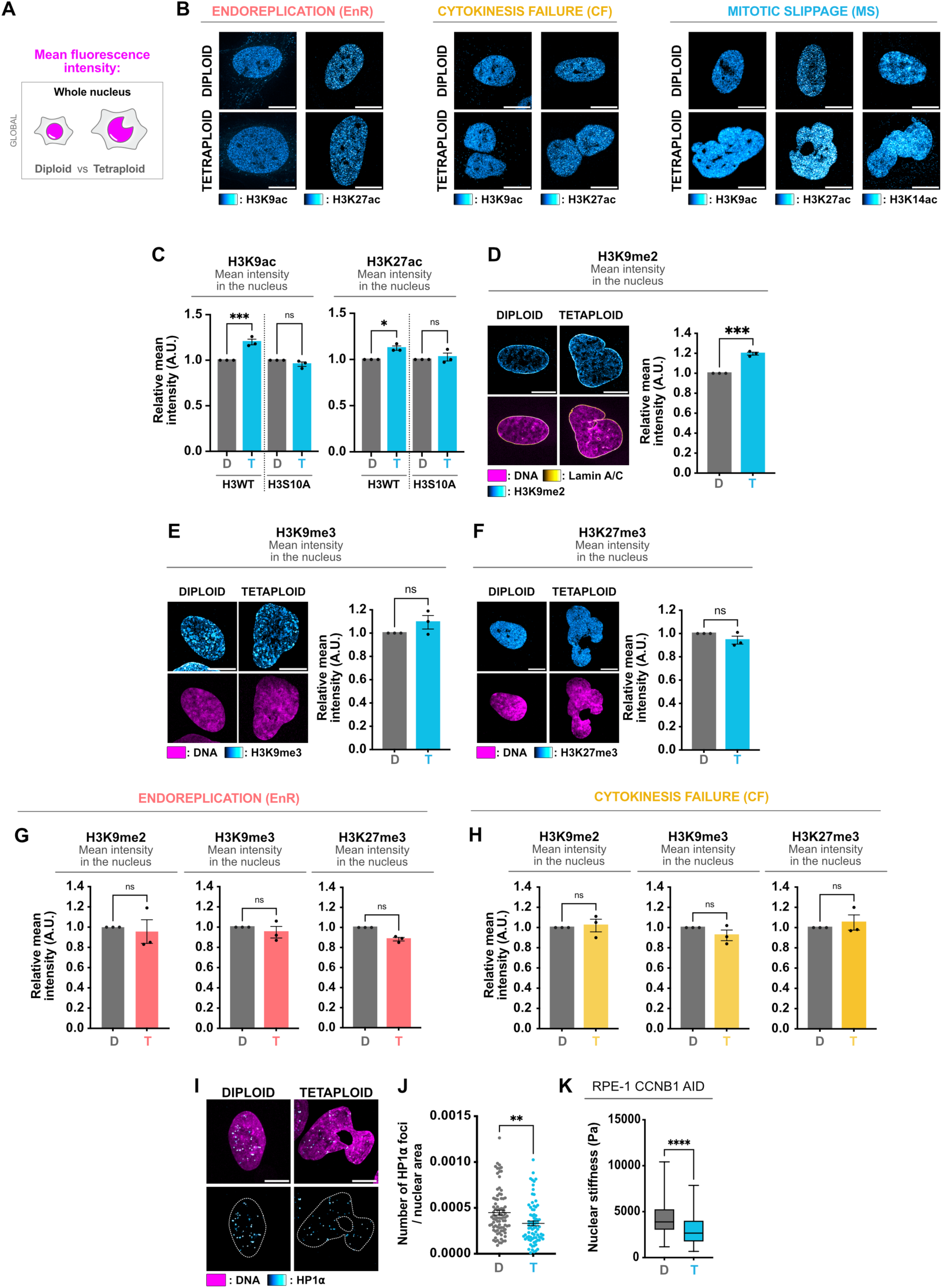
Mitotic slippage, but not cytokinesis failure and endoreplication, alters histone modification **(A)** Schematic representation of the mean fluorescence measured at the level of the whole nucleus in diploid and tetraploid cells. **(B)** SIM representative images showing diploid and tetraploid G1 RPE-1 cells generated by EnR (left panel), by CF (central panel) or by MS (right panel). H3K9ac, H3K27ac and H3K14ac in cyan hot. **(C)** Graphs showing the relative H3K9ac (left panel) or H3K27ac (right panel) mean intensity in the nucleus in diploid (in grey) and in tetraploid (in blue) RPE-1 G1 cells stably expressing H3 WT or H3 S10A. Mean ± SEM, >100 G1 cells analysed per condition, three independent experiments. **(D)** Left panel – SIM representative images showing diploid and tetraploid G1 RPE-1 cells generated by MS. Lamin A/C in yellow hot, H3K9me2 in cyan hot and DNA in magenta. Right panel – Graph presenting the relative H3K9me2 mean intensity in the nucleus in diploid (in grey) and in tetraploid (in blue) G1 RPE-1 cells. Mean ± SEM, >100 G1 cells analysed per condition, three independent experiments. **(E)** Left panel - SIM representative images showing diploid and tetraploid G1 RPE-1 cells generated by MS. Lamin A/C in yellow hot, H3K9me3 in cyan hot and DNA in magenta. Right panel – Graph presenting the relative H3K9me3 mean intensity in the nucleus in RPE-1 diploid (in grey) and in tetraploid (in blue) G1 cells. Mean ± SEM, >100 G1 cells analysed per condition, three independent experiments. **(F)** Left panel - SIM representative images showing diploid and tetraploid G1 RPE-1 cells generated by MS. Lamin A/C in yellow hot, H3K27me3 in cyan hot and DNA in magenta. Right panel – Graph presenting the Relative H3K27me2 mean intensity in the nucleus in diploid (in grey) and in tetraploid (in blue) G1 RPE-1 cells. Mean ± SEM, >100 G1 cells analysed per condition, three independent experiments. **(G-H)** Graphs showing the relative H3K9me2, H3K9me3 and H3K27me3 mean intensity in the nucleus in diploid (in grey) and in tetraploid G1 RPE-1 cells generated by **(G)** EnR (in red) or by **(H)** CF (in yellow). Mean ± SEM, >100 G1 cells analysed per condition, three independent experiments. **(I)** SIM representative images showing diploid and tetraploid G1 RPE-1 cells generated by MS. HP1α in cyan hot and DNA in magenta. **(J)** Graph showing the number of HP1 foci / nuclear area in diploid (in grey) and in tetraploid (in blue) RPE-1 G1 cells. Mean ± SEM, >90 G1 cells analysed per condition, at least three independent experiments. **(K)** Graph showing nuclear stiffness in diploid and in MS-generated tetraploid G1 RPE-1 CCNB1 cells (in grey and blue, respectively). Box and whiskers ± Min to Max, >175 G1 cells analysed per condition, three independent experiments. Scale bars: 10µm. D=diploid. T=tetraploid. MS=mitotic slippage. EnR=endoreplication. CF=cytokinesis failure. A.U.=arbitrary unit. **(C,D,E,F,G,H,J)** t-test (two-sided). **(K)** Kolmogorov-Smirnov test. ns=not significant. *=P ≤ 0.05. **= P ≤ 0.01. ***= P ≤ 0.001. ****= P ≤ 0.0001.

**Figure S4:**
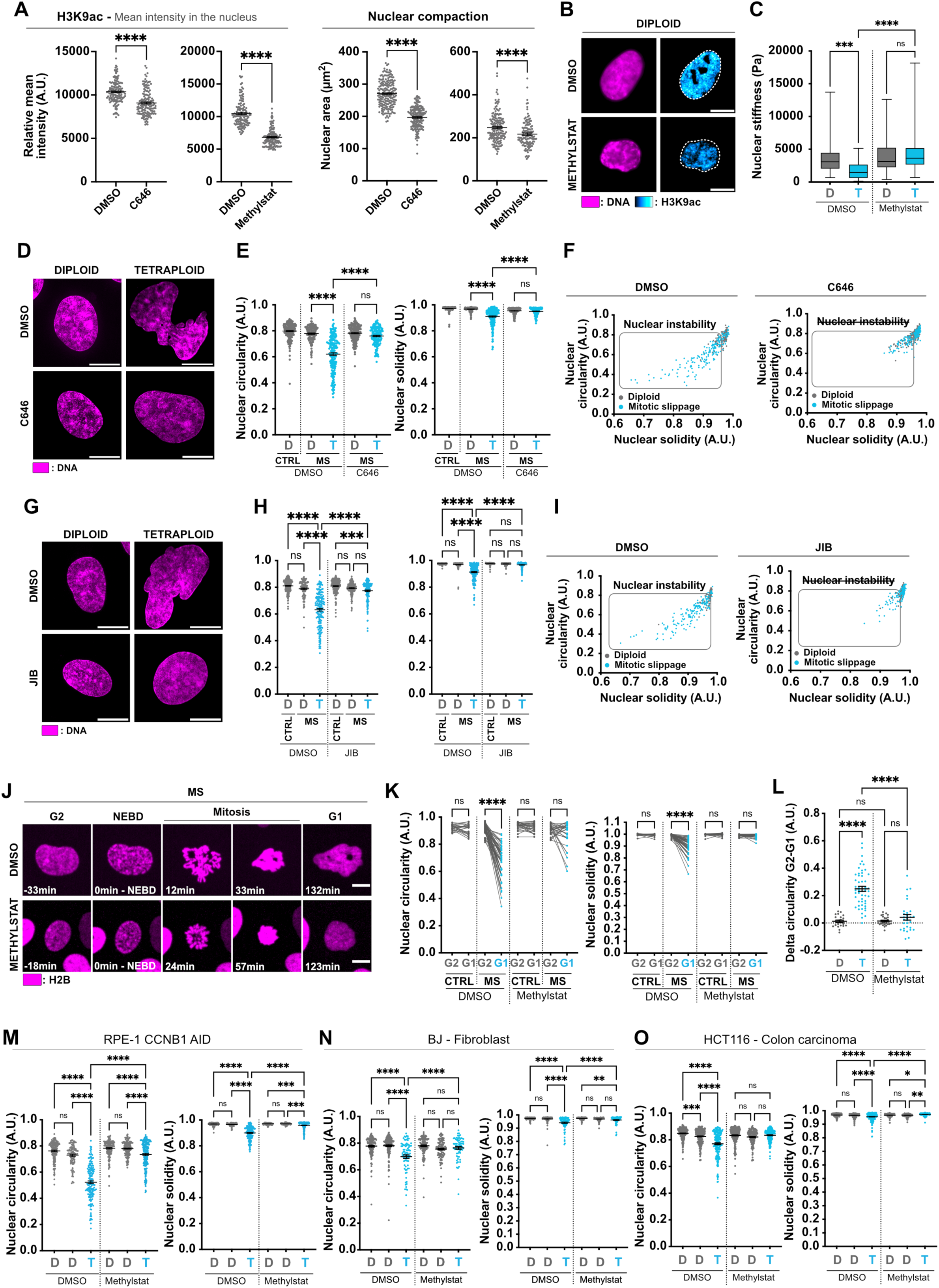
Promoting nuclear compaction is sufficient to prevent nuclear deformations upon mitotic slippage **(A)** Graphs presenting the relative H3K9ac mean intensity (left panel) or the nuclear area (right panel) in diploid RPE-1 cells. Mean ± SEM, >125 cells analysed per condition, at least three independent experiments. **(B)** Representative images showing RPE-1 diploid cells treated with DMSO or Methylstat. H3K9ac in cyan hot, DNA in magenta. **(C)** Graph showing the nuclear stiffness in diploid (in grey) and tetraploid (in blue) RPE-1 G1 cells generated by MS treated with DMSO or Methylstat. **(D)** Representative images showing diploid and tetraploid RPE-1 G1 cells treated with DMSO or C646. DNA in magenta. **(E)** Graphs showing the nuclear circularity (left panel) and solidity (ritght panel) indicies in diploid (in grey) and tetraploid (in blue) RPE-1 G1 cells generated by MS. Mean ± SEM, >125 G1 cells analysed per condition, at least three independent experiments. **(F)** Graphs presenting the nuclear circularity and solidity from **(E)** in RPE-1 diploid (in grey) and tetraploid (in blue) G1 cells treated with DMSO or C646. **(G)** Representative images showing diploid and tetraploid RPE-1 cells treated with DMSO or JIB. DNA in magenta. **(H)** Graphs presenting the nuclear circularity (left panel) and solidity (ritght panel) indices in diploid (in grey) and tetraploid (in blue) RPE-1 G1 cells generated by MS treated with DMSO or 5µM JIB. Mean ± SEM, >100 G1 cells analysed per condition, at least three independent experiments. **(I)** Graphs presenting the nuclear circularity and solidity from **(H)** in diploid (in grey) and tetraploid (in blue) RPE-1 G1 cells. **(J)** Stills from time-lapse imaging of RPE-1 cells stably expressing mCherry-H2B (in magenta) performing MS and treated with DMSO (upper panel) or Methylstat (lower panel). **(K)** Graphs showing the nuclear circularity (left panel) and solidity (right panel) measured at the single cell levels in G2 and in G1 upon mitotic division (in grey) or MS (in blue) in RPE-1 cells treated with DMSO or 2µM Methylstat. >30 cells analysed per condition, three independent experiments. **(L)** Graph showing the delta circularity (defined as the difference between nuclear circularity in G2 vs in G1) upon mitotic division (in grey) or MS (in blue) in RPE-1 cells treated with DMSO or 2µM Methylstat. Mean ± SEM, >25 cells analysed per condition, three independent experiments. **(M-O)** Graphs displaying nuclear circularity and solidity in **(M)** RPE-CCNB1 or **(N)** BJ or **(O)** HCT116 cells treated with DMSO or Methylstat. Mean ± SEM, >65 G1 cells analysed per condition, three independent experiments. Scale bars: 10µm. D=diploid. T=tetraploid. MS=mitotic slippage. NEBD=nuclear envelope breakdown. A.U.=arbitrary unit**. (A,K,L)** t-test (two-sided). **(E,H,M,N,O)** Anova-test (one-sided). **(C)** Kolmogorov-Smirnov test. ns=not significant. *=P ≤ 0.05. **= P ≤ 0.01. ***= P ≤ 0.001. ****= P ≤ 0.0001.

**Figure S5:**
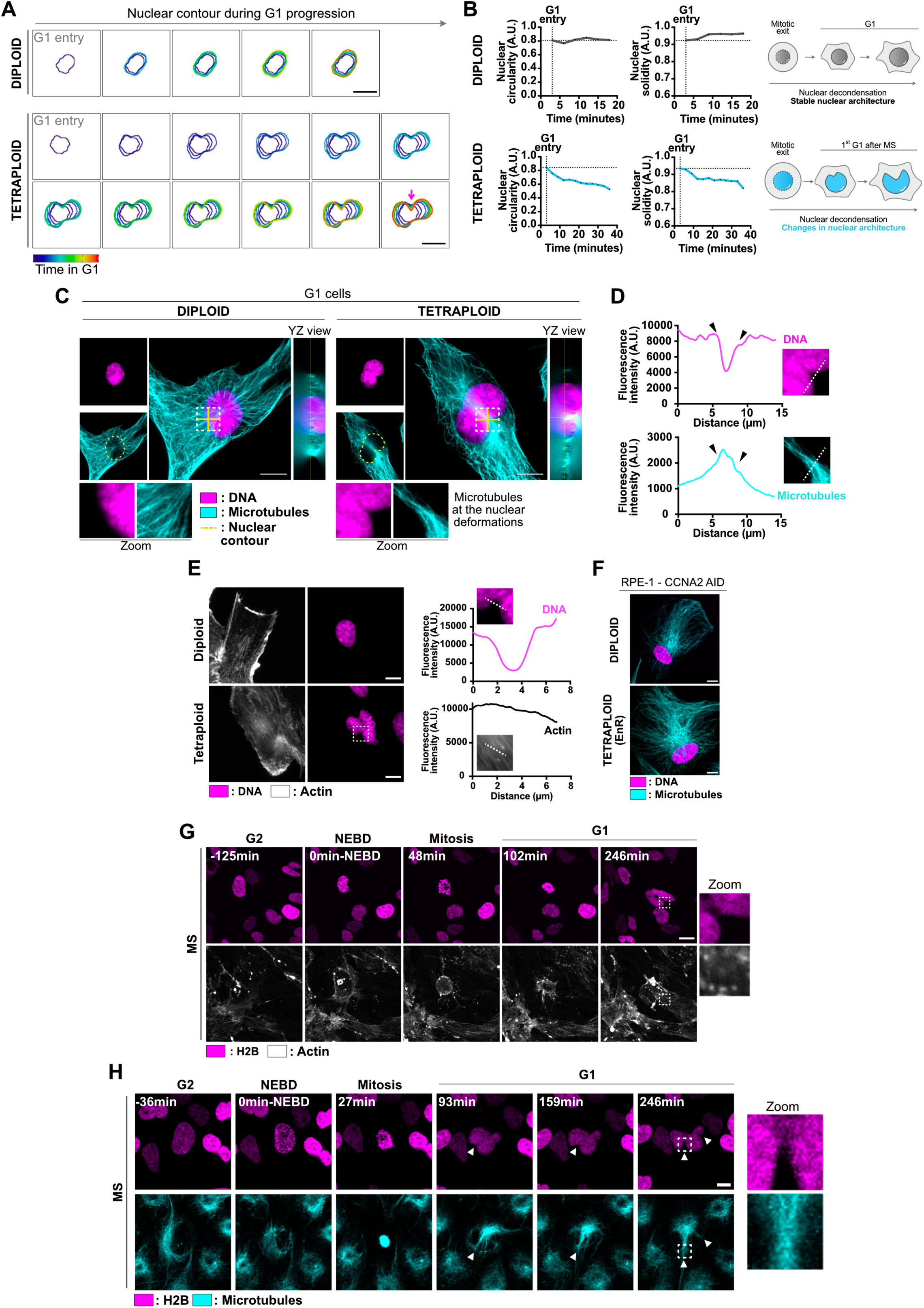
The microtubule cytoskeleton contributes to nuclear deformations in mitotic slippage-induced newly born tetraploid cells. (**A)** Stills from time-lapse imaging of RPE-1 cells stably expressing mCherry-H2B after mitotic division (upper panel) or MS (lower panel). The nuclear contour during G1 is indicated, and the colour code shows the time spent in G1 (see methods). The pink arrow points at a nuclear deformation. **(B)** Left panel - Graph showing the nuclear circularity and solidity in G1 upon mitotic division (in grey, upper panel) or MS (in blue, lower panel). Right panel - Schematic representations of nuclear decondensation upon mitotic division (in grey, upper panel) or MS (in blue, lower panel). The data were measured in the cells presented in **(A**). **(C)** Representative images of diploid and MS-generated tetraploid RPE-1 G1 cells. Microtubules in cyan, DNA in magenta. The nuclear contour is indicated (yellow dotted line). **(D)** Graphs showing line scans measurements. **(E)** Left panel - Representative images showing diploid and MS-generated RPE-1 cells. Actin in grey, DNA in magenta. Right panel - Graphs showing line scan measurements. **(F)** Representative images showing microtubules (in cyan) in diploid and in EnR-generated RPE-1 cells. DNA in magenta. **(G)** Stills from time-lapse imaging of RPE-1 cells stably expressing mCherry-H2B (in magenta) performing MS. Actin was labelled using SPY650-FastAct (in grey). **(H)** Time-lapse still images from movies of RPE-1 cells stably expressing mCherry-H2B (in magenta undergoing MS. Microtubules in cyan. Scale bars: 10µm. D=diploid. T=tetraploid. MS=mitotic slippage. NEBD=nuclear envelope breakdown. A.U.=arbitrary unit.

**Figure S6:**
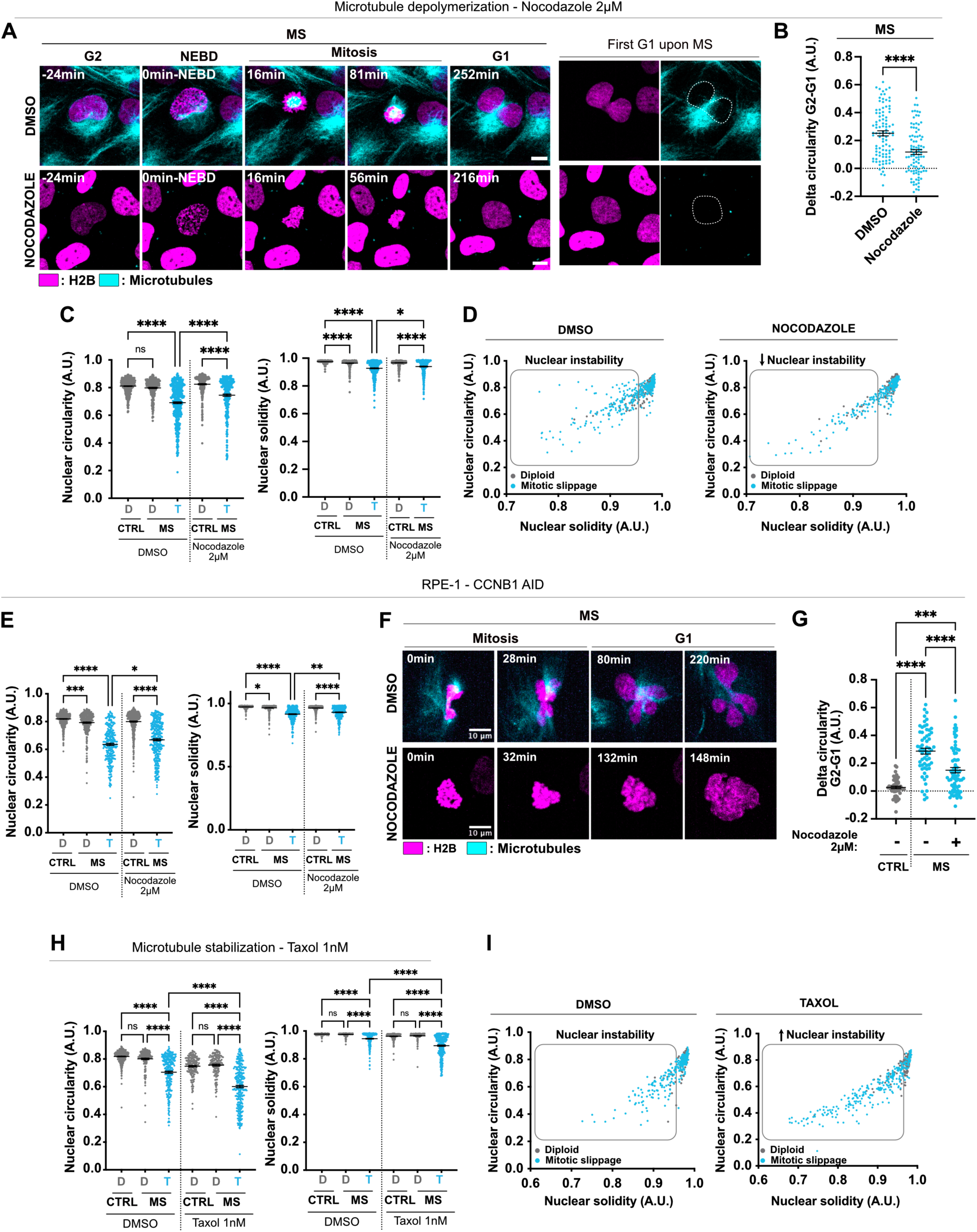
The microtubule cytoskeleton is the main contributor to nuclear deformations in tetraploid cells generated by mitotic slippage. **(A)** Stills from time-lapse imaging of RPE-1 cells stably expressing mCherry-H2B (in magenta) undergoing MS and treated with DMSO (upper panel) or nocodazole (lower panel). Microtubules in cyan. **(B)** Graph presenting the delta circularity (defined as the difference between nuclear circularity in G2 vs in G1) in MS-generated tetraploid cells treated with DMSO or Nocodazole. Mean ± SEM, >90 cells analysed per condition, three independent experiments. **(C)** Graphs showing nuclear circularity (left panel) and solidity (right panel) in diploid and in MS-generated tetraploid RPE-1 cells (in grey and blue, respectively) treated with DMSO or with 2µM Nocodazole. Mean ± SEM, >105 G1 cells analysed per condition, three independent experiments. **(D)** Graphs presenting the nuclear circularity and solidity from **(C)** in diploid and tetraploid G1 RPE-1 cells treated with DMSO or with Nocodazole. **(E)** Graphs showing nuclear circularity and solidity in diploid and in MS-generated tetraploid RPE-1 CCNB1 cells (in grey and blue, respectively) treated with DMSO or with 2µM Nocodazole. Mean ± SEM, >80 G1 cells analysed per condition, three independent experiments. **(F)** Stills from time-lapse imaging of RPE-1 CCNB1 cells stably expressing mCherry-H2B (in magenta) treated with DMSO or Nocodazole and performing MS. **(G)** Graph showing the delta circularity (defined as the difference between nuclear circularity in G2 vs in G1) upon mitotic division (in grey) or MS (in blue) treated with DMSO (-) or Nocodazole. Mean ± SEM, >50 cells analysed per condition, three independent experiments. **(H)** Graphs displaying nuclear circularity and solidity indices in diploid and in MS-generated tetraploid RPE-1 cells (in grey and blue, respectively) treated with DMSO or Taxol. Mean ± SEM, >140 G1 cells analysed per condition, three independent experiments. **(I)** Graph presenting the nuclear circularity and solidity from **(H)** in diploid and tetraploid G1 RPE-1 cells treated with DMSO or with 1nM Taxol. Scale bars: 10µm. D=diploid. T=tetraploid. MS=mitotic slippage. CF=cytokinesis failure. EnR= endoreplication. A.U.=arbitrary unit. **(B)** t-test (two-sided). **(C,E,G,H)** Anova-test (one-sided). ns=not significant. *=P ≤ 0.05. **= P ≤ 0.01. ***= P ≤ 0.001. ****= P ≤ 0.0001.

**Figure S7:**
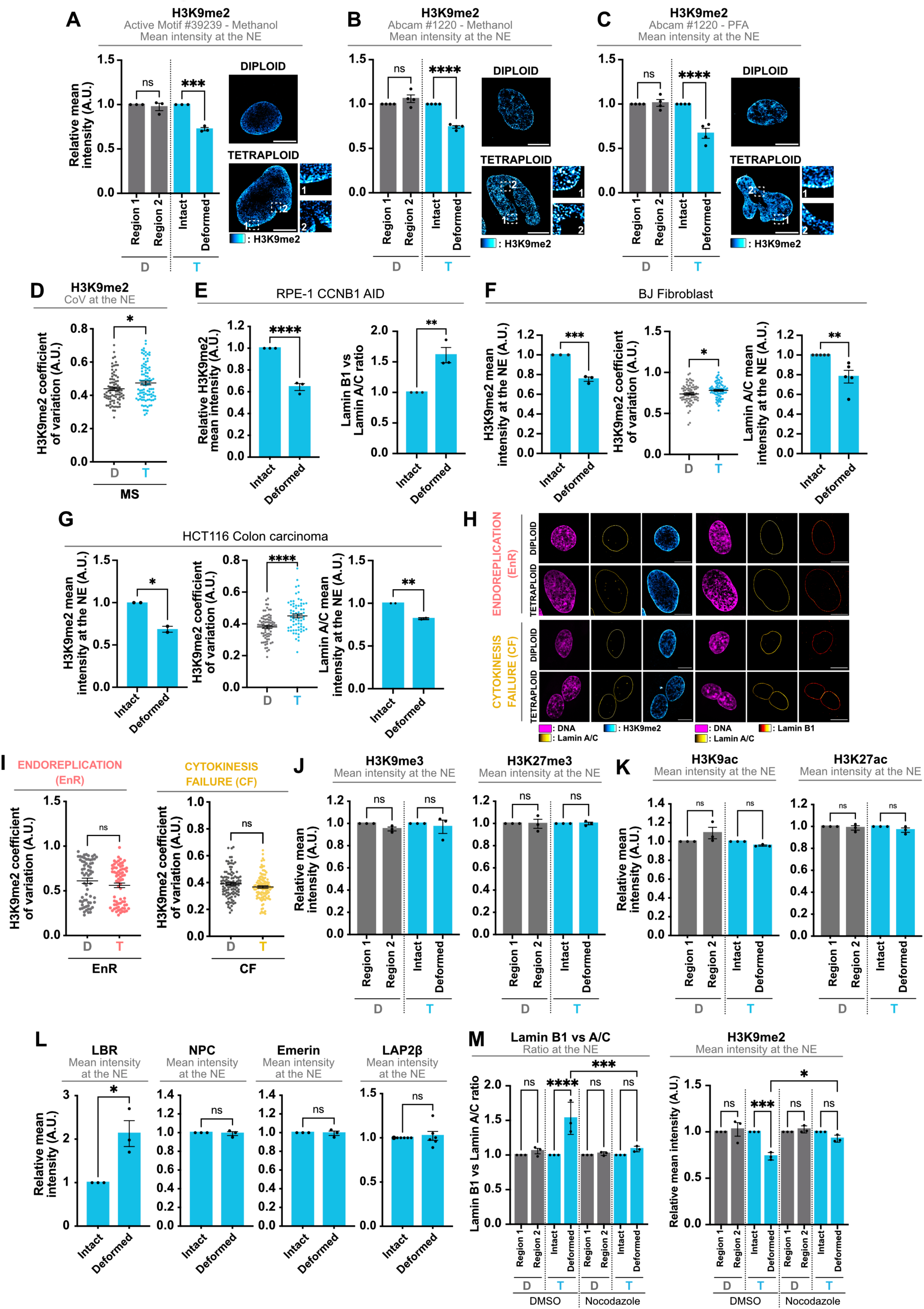
Mitotic slippage promotes local changes in the Lamina network and in H3K9me2 levels **(A)** Left panel – Graph showing the relative H3K9me2 mean intensity at the nuclear envelope in diploid (in grey) and tetraploid (in blue) G1 RPE-1 cells. Right panel – SIM representative images showing diploid and tetraploid G1 RPE-1 cells. H3K9me2 in cyan hot. Mean ± SEM, >90 G1 cells analysed per condition, three independent experiments. **(B)** Left panel – Graph showing the relative H3K9me2 mean intensity at the nuclear envelope in diploid (in grey) and tetraploid (in blue) G1 RPE-1 cells. Mean ± SEM, >90 G1 cells analysed per condition, three independent experiments. Right panel – SIM representative images showing diploid and tetraploid G1 RPE-1 cells. H3K9me2 in cyan hot. **(C)** Left panel – Graph showing the relative H3K9me2 mean intensity at the nuclear envelope in diploid (in grey) and tetraploid (in blue) G1 RPE-1 cells. Mean ± SEM, >90 G1 cells SIM resolution representative images showing diploid and tetraploid G1 RPE-1 cells. H3K9me2 in cyan hot. **(D)** Graph presenting the H3K9me2 coefficient of variation at the nuclear envelope in MS-induced tetraploid G1 RPE-1 cells. Mean ± SEM, >90 G1 cells analysed per condition, three independent experiments. **(E)** Graphs showing the relative H3K9me2 mean intensity (left panel) or the Lamin B1 vs A/C ratio (right panel) at the nuclear envelope in MS-induced tetraploid RPE-1 CCNB1 cells. Mean ± SEM, >90 G1 cells analysed per condition, three independent experiments. **(F-G)** Graphs of H3K9me2 mean intensity at the NE (left panel), H3K9me2 coefficient of variation at the NE (central panel) and Lamin A/C mean intensity at the NE (right panel) in diploid and in MS-generated tetraploid **(F)** BJ and **(G)** HCT116 cells. Mean ± SEM, >60 G1 cells analysed per condition, three independent experiments. **(H)** SIM representative images showing diploid and tetraploid cells generated by EnR or CF. H3K9me2, Lamin A/C and Lamin B1 in cyan hot, yellow hot and red hot, respectively. DNA in magenta. **(I)** Graphs showing the H3K9me2 coefficient of variation in diploid (in grey) and tetraploid G1 RPE-1 cells generated by EnR (in red) or CF (in yellow). Mean ± SEM, >90 G1 cells analysed per condition, three independent experiments. **(J)** Graphs showing the relative H3K9me3 (left panel) or H3K27me3 (right panel) mean intensity in diploid (in grey) and tetraploid (in blue) RPE-1 G1 cells. Mean ± SEM, >90 G1 cells analysed per condition, three independent experiments. **(K)** Graphs showing the relative H3K9ac (left panel) or H3K27ac (right panel) mean intensity in diploid (in grey) and tetraploid (in blue) RPE-1 G1 cells. Mean ± SEM, >90 G1 cells analysed per condition, three independent experiments. **(L)** Graphs presenting the relative mean intensity at the nuclear envelope of the indicated proteins in tetraploid RPE-1 G1 cells. Mean ± SEM, >90 G1 cells analysed per condition, at least three independent experiments. **(M)** Graphs showing the Lamin B1 vs A/C ratio (left panel) or the H3K9me2 mean intensity (right panel) at the nuclear envelope in diploid (in grey) and tetraploid (in blue) RPE-1 G1 cells treated with DMSO or with Nocodazole. Mean ± SEM, >90 G1 cells analysed per condition, three independent experiments. Scale bars: 10µm. D=diploid. T=tetraploid. MS=mitotic slippage. EnR=Endoreplication. CF=Cytokinesis failure. A.U.=arbitrary unit. **(M)** Anova-test (one-sided). **(A,B,C,D,E,F,G,I,J,K,L)** t-test (two-sided). ns=not significant. *=P ≤ 0.05. **= P ≤ 0.01. ***= P ≤ 0.001. ****= P ≤ 0.0001.

**Figure S8:**
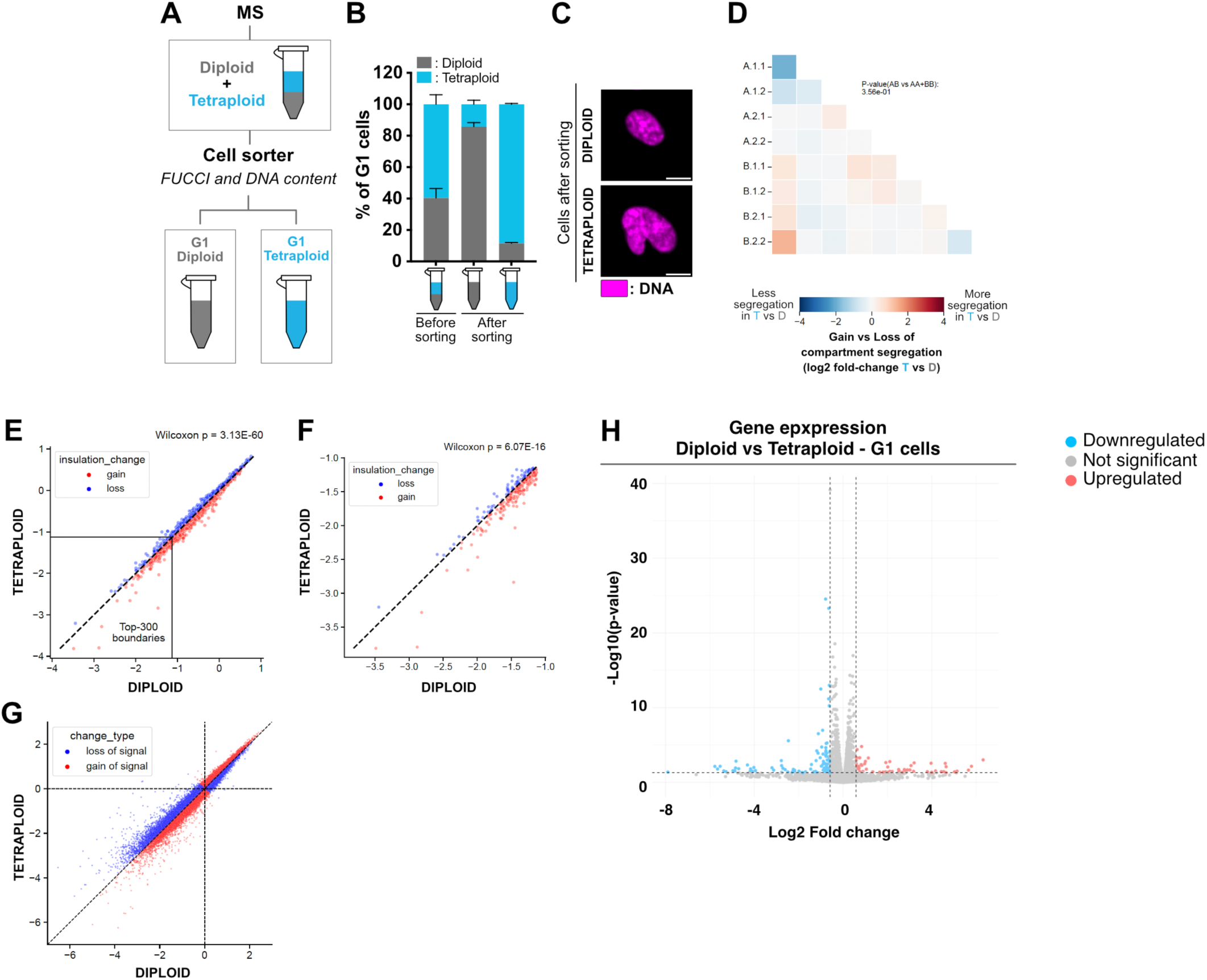
Comparison of chromosome interactions and gene expression in tetraploid and diploid RPE1 cells **(A)** Scheme presenting the experimental workflow used to isolate diploid and tetraploid cells. **(B)** Graph showing the percentage of G1 diploid (in grey) and MS-generated tetraploid (in blue) RPE-1 cells before and after cell sorting. Mean ± SEM, >900 cells analysed, five independent experiments. **(C)** Representative images showing diploid and MS-generated tetraploid cells after cell sorting. DNA in magenta. **(D-E)** Boundary insulation scores for **(D)** all and **(E)** top 300 most strongly insulating boundaries between diploid (x-axis) and tetraploid (y-axis) conditions. **(F)** Relationship between observed and expected intra-chromosomal values in tetraploid versus diploid. Each dot represents an inter-chromosomal interaction, red signifies higher and blue lower interaction signal in tetraploid compared to control. **(G)** Remodeling of interactions between chromatin sub-compartments in tetraploid and diploid cells. Blue suggests a loss of interaction, red a gain of interactions. **(H)** Volcano plot of RNAseq data obtained by comparing gene expression in tetraploid G1 cells generated by MS with diploid G1 cells. D=diploid. T=tetraploid. MS=mitotic slippage. ns=not significant. **(D,E,F,G)** two-tailed Wilcoxon test. *=P ≤ 0.05. **= P ≤ 0.01. ***= P ≤ 0.001. ****= P ≤ 0.0001.

**Figure S9:**
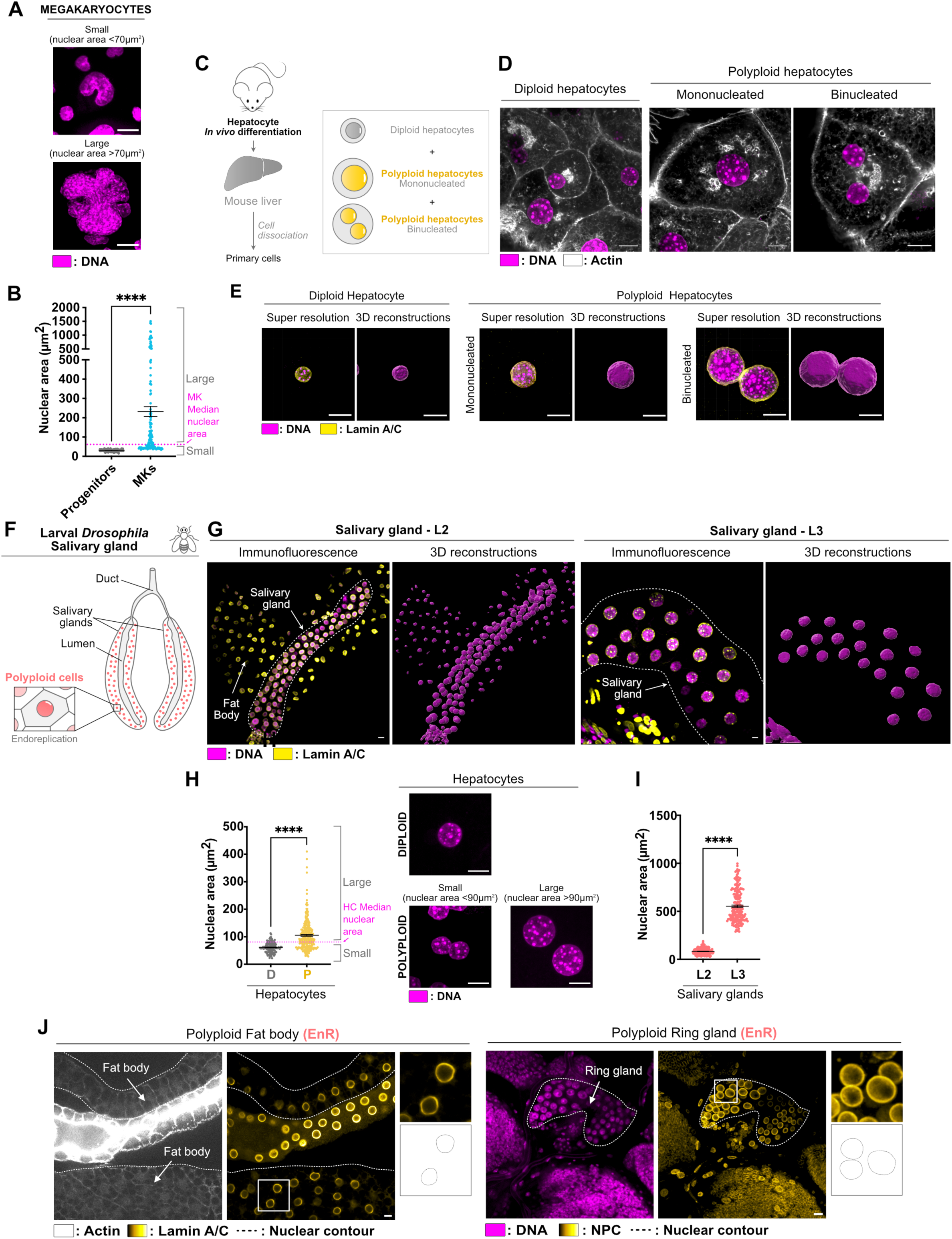
Physiological polyploid cells generated by cytokinesis failure and endoreplication show homogenous nuclear shape **(A)** Representative images of small and large MKs. DNA in magenta. **(B)** Graph showing nuclear area in diploid progenitors and in polyploid MKs. The MK median nuclear area was used to distinguish small and large MKs. Mean ± SEM, >150 interphase cells analysed per condition, three independent experiments. **(C)** Schematic workflow showing the methods used to obtain mouse hepatocytes. **(D)** Representative images showing diploid and polyploid hepatocytes. Actin in grey, DNA in magenta. **(E)** SIM representative images and their corresponding 3D reconstructions showing diploid and polyploid hepatocytes. Lamin A/C in yellow, DNA in magenta. **(F)** Schematic representation of *Drosophila* salivary glands. **(G)** Representative images and their corresponding 3D reconstructions showing polyploid salivary glands. Lamin A/C in yellow, DNA in magenta. **(H)** Left panel - Graph showing the nuclear area in diploid and in polyploid hepatocytes (in grey and yellow, respectively). The hepatocyte median nuclear area was used to distinguish small and large hepatocytes. Mean ± SEM, >140 interphase cells analysed per condition, three independent experiments. Right panel – Representative images showing diploid and polyploid hepatocytes. DNA in magenta. **(I)** Graph showing the nuclear area and in polyploid salivary glands at two different developmental stages (L2 and L3). Mean ± SEM, >100 cells analysed, three independent experiments. **(J)** Representative immunofluorescence images showing *Drosophila* fat body (left panel) and ring gland (right panel). Lamin A/C or Nuclear pore complex (NPC) in yellow hot, Actin in grey, DNA in magenta. Scale bars: 10µm. D=diploid. P=polyploid. MK=megakaryocyte. NPC=Nuclear pore complex. **(B,H,I)** t-test (two-sided). ns=not significant. *=P ≤ 0.05. **= P ≤ 0.01. ***= P ≤ 0.001. ****= P ≤ 0.0001.

**Figure S10:**
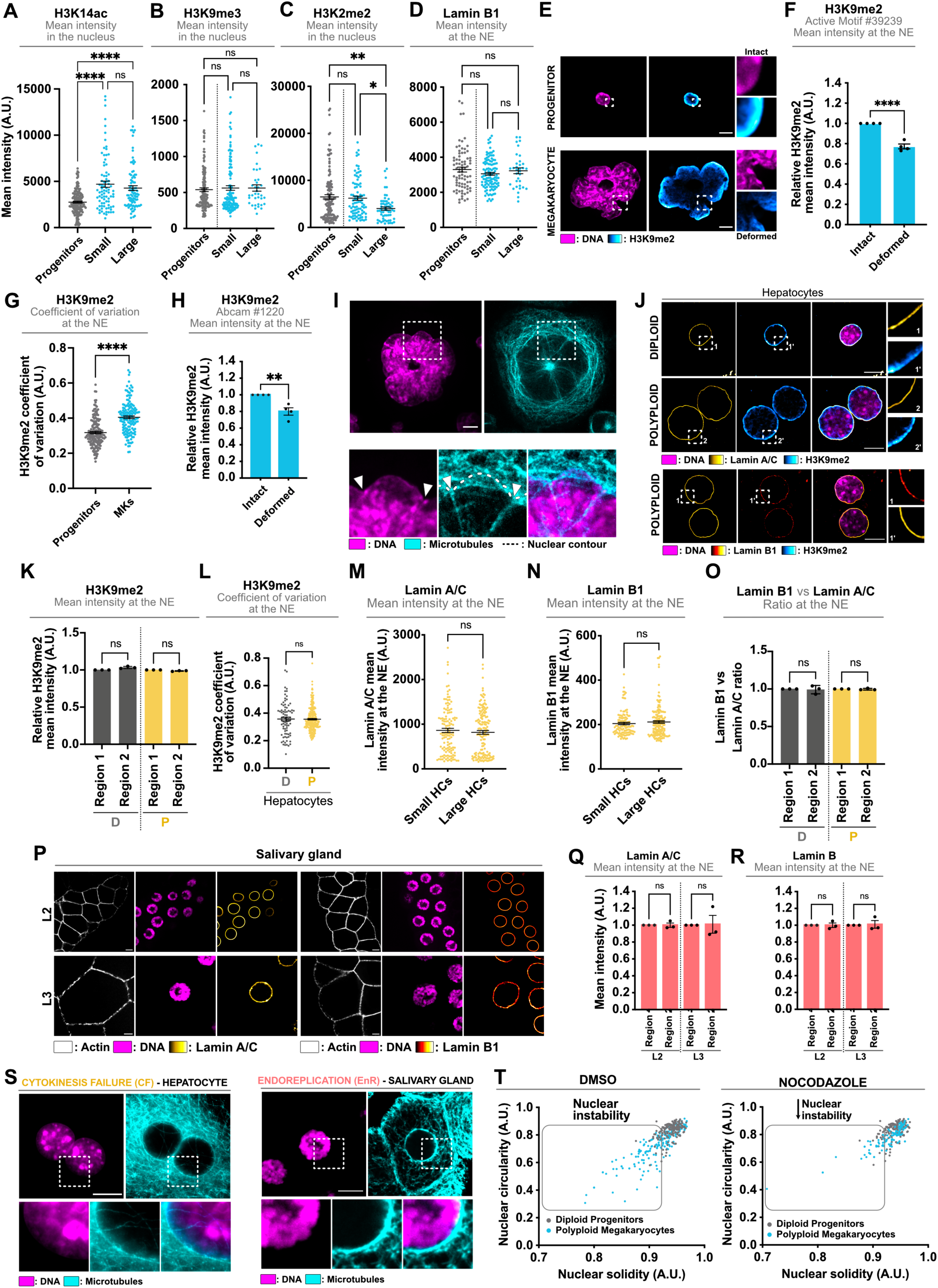
Polyploid hepatocytes and salivary gland cells do no show defects in H3K9me2 and Lamin A/C distribution **(A-D)** Graphs showing **(A)** H3K27ac, **(B)** H3K9me3 and **(C)** H3K9me2 mean intensity in the nucleus in diploid progenitors (in grey) and in polyploid megakaryocytes (in blue). Mean ± SEM, >120 cells analysed, three independent experiments and Lamin B1 mean intensity at the nuclear envelope **(D)**in diploid progenitors (in grey) and in polyploid megakaryocytes (in blue). **(E)** SIM representative images of diploid progenitors and polyploid megakaryocyte. H3K9me2 in cyan hot, DNA in magenta. **(F-G)** Graphs showing H3K9me2 **(F)** mean intensity and **(G**) coefficient of variation (define as the ratio of the standard deviation to the mean) in progenitors (in grey) and in MKs (in blue). Mean ± SEM, >150 interphase cells analysed per condition, four independent experiments. **(H)** Graph showing H3K9me2 mean intensity in progenitors (in grey) and in MKs (in blue). Mean ± SEM, >65 interphase cells analysed per condition, four independent experiments. **(I)** Representative images of microtubules (in cyan) in MKs, DNA in magenta. **(J)** SIM representative images showing diploid and polyploid hepatocytes. Lamin A/C, Lamin B1 and H3K9me2 in yellow hot, red hot and cyan hot, respectively, DNA in magenta. **(K-L)** Graphs showing **(K)** the relative H3K9me2 mean intensity at the NE and **(L)** the H3K9me2 coefficient of variation (define as the ratio of the standard deviation to the mean) at the NE in diploid (in grey) and in polyploid (in yellow) hepatocytes. Mean ± SEM, >90 interphase cells analysed per condition, three independent experiments. **(M-O)** Graphs presenting **(M)** Lamin A/C and **(N**) Lamin B1 mean intensity at the NE and **(O)** the Lamin B1 vs Lamin A/C ratio at the NE in hepatocytes. Mean ± SEM, >90 interphase cells analysed per condition, three independent experiments. **(P)** Representative images showing *Drosophila* salivary glands at two different developmental stages (L2 and L3). Lamin A/C and Lamin B1 in yellow hot and red hot, respectively, Actin in grey, DNA in magenta. **(Q-R)** Graphs showing **(Q)** Lamin A/C and **(R)** Lamin B1 mean intensity at the NE in *Drosophila* salivary glands. Mean ± SEM, >150 cells analysed per condition, three independent experiments. **(S)** Representative images showing (left panel) polyploid hepatocytes and (right panel) *Drosophila* salivary gland cells. Microtubules in cyan. DNA in magenta. **(T)** Graph presenting the nuclear circularity and solidity index in progenitors (in grey) or megakaryocytes (in blue) treated with DMSO or Nocodazole. Scale bars: 10µm. D=diploid. P=polyploid. HC=hepatocytes. MK=megakaryocytes. NE=nuclear envelope. **(D)** Anova-test (one-sided). **(F,G,J,K,L,M,N,P)** t-test (two-sided). ns=not significant. *=P ≤ 0.05. **= P ≤ 0.01. ***= P ≤ 0.001. ****= P ≤ 0.0001. Videos 1 and 2: Mitotic slippage generates tetraploid cells with nuclear deformations. Time lapse videos of RPE-1 cells expressing mCherry-H2B (in magenta) and performing canonical mitosis (video 1) or mitotic slippage (video 2). Images were acquired every 5 minutes. Videos 3 and 4: Microtubules accumulate around nuclear deformations upon mitotic slippage. Time lapse videos of RPE-1 cells expressing mCherry-H2B (in magenta) performing mitotic slippage. Microtubules were labeled using SPY Tubulin (in Cyan, video 3) and actin was labeled using SPY FAST Actin (in blue, video 4). Images were acquired every 3 minutes. Videos 5 and 6: Microtubule depolymerization prevents nuclear deformations upon mitotic slippage. Time lapse videos of RPE-1 cells expressing mCherry-H2B (in magenta) and performing mitotic slippage in the absence (video 5) or presence of Nocodazole (video 6). Microtubules were labeled using SPY Tubulin (in Cyan). Images were acquired every 3 minutes. Videos 7 and 8: Microtubule depolymerization prevents nuclear deformations upon mitotic slippage induced by CCNB1 depletion. Time lapse videos of RPE-1 CCNB1 cells expressing mCherry-H2B (in magenta) and performing mitotic slippage in the absence (video 7) or presence of Nocodazole (video 8). Microtubules were labeled using SPY Tubulin (in Cyan). Images were acquired every 3 minutes. Videos 9 and 10: Megakaryocytes become polyploid by mitotic slippage. Time lapse videos Megakaryocytes expressing mCherry-H2B (in magenta) and performing mitotic slippage. Images were acquired every 10 minutes.

